# FAM172A controls the nuclear import and alternative splicing function of AGO2

**DOI:** 10.1101/2022.03.04.482948

**Authors:** Sephora Sallis, Félix-Antoine Bérubé-Simard, Benoit Grondin, Elizabeth Leduc, Fatiha Azouz, Catherine Bélanger, Nicolas Pilon

## Abstract

The poorly characterized protein FAM172A is mutated in some individuals affected by a disorder of neural crest development called CHARGE syndrome. We also know that FAM172A can interact with the main CHARGE syndrome-associated protein CHD7 and the small RNA-binding protein AGO2 at the chromatin-spliceosome interface. Focusing on this intriguing FAM172A-AGO2 interaction, we now report that FAM172A is one of the long sought-after regulator of AGO2 nuclear import. This FAM172A function relies on its nuclear localization signal, being enhanced by CK2-mediated phosphorylation and abrogated by a CHARGE syndrome-associated missense mutation. Accordingly, *Fam172a* and *Ago2* genetically interact in mice, and neural crest-specific depletion of *Ago2* is sufficient to phenocopy CHARGE syndrome without impacting post-transcriptional gene silencing. Rapamycin-mediated rescue suggests that observed morphological anomalies are instead due to alternative splicing defects. This work thus demonstrates that non-canonical nuclear functions of AGO2 and associated regulatory mechanisms may be clinically relevant.

## Introduction

CHARGE syndrome is a severe multi-organ developmental disorder mainly affecting derivatives of cranial and cardiac neural crest cells (NCCs) (Pauli, Bajpai et al., 2017, Pilon, 2021). Many, but not all, of these anomalies are included in the CHARGE acronym: <u>C</u>oloboma of the eye, <u>H</u>eart defects, <u>A</u>tresia of choanae, <u>R</u>etardation of growth/development, <u>G</u>enital abnormalities, and <u>E</u>ar anomalies. Other frequently encountered anomalies include craniofacial dysmorphism and cleft palate (Hale et al., 2016). Heterozygous mutation of *CHD7* (*Chromodomain Helicase DNA Binding Protein-7*) is the main cause of CHARGE syndrome (Vissers et al., 2004). Yet, rare variants in many other genes encoding chromatin and/or splicing factors have also been recently associated with this disorder, including *PUF60, EP300, RERE, KMT2D, KDM6A* and *FAM172A* (Bélanger et al., 2018; Moccia et al., 2018).

We previously validated the candidacy of *FAM172A* (*Family With Sequence Similarity 172 Member A*) as CHARGE syndrome-associated gene using a mouse model issued from a forward genetic screen (Pilon, 2016). In this model called *Toupee*, mutagenic transgene sequences are inserted in the last intron of *Fam172a* (Belanger, Berube-Simard et al., 2018), generating a hypomorphic allele that negatively affects almost every aspect of NCC ontology (Belanger et al., 2018). Accordingly, homozygous *Toupee* animals (*Fam172a*^*Tp/Tp*^) phenocopy both “major” (*e*.*g*., coloboma, cleft palate and vestibular hypoplasia) and “minor” (*e*.*g*., retarded growth, cardiac malformations, cranial nerve defects and genital anomalies) features of CHARGE syndrome (Belanger et al., 2018, Hale, Niederriter et al., 2016). Moreover, the *Toupee* allele was found to genetically interact with a gene-trap mutant allele of the main CHARGE syndrome-associated gene *Chd7* (Belanger et al., 2018, Hurd, Capers et al., 2007). Our mechanistic studies further allowed us to propose a model whereby FAM172A – which contains a large ARB2 domain (Argonaute Binding protein 2) split in two halves by a bipartite nuclear localization signal (NLS) – helps to stabilize the chromatin-spliceosome interface, notably by interacting with CHD7 and AGO2 (Belanger et al., 2018). These studies also suggested that perturbation of chromatin-mediated alternative splicing is a general pathological mechanism for CHARGE syndrome, regardless of involved gene defect (Belanger et al., 2018, Belanger, Cardinal et al., 2022, Berube-Simard & Pilon, 2018).

AGO2 (Argonaute-2) is a small RNA-binding protein best known as major component of the RNA-induced silencing complex (RISC), which orchestrates post-transcriptional gene silencing in the cytoplasm (Hutvagner and Simard, 2008; Liu et al., 2004). Yet, increasing evidence suggests that AGO2 also fulfills important functions in the nucleus (Nazer, Gomez Acuna et al., 2022), notably for regulating alternative splicing (Ameyar-Zazoua, Rachez et al., 2012, Batsche & Ameyar-Zazoua, 2015, Chu, Yokota et al., 2021, Kalantari, Chiang et al., 2016, Meng, Ma et al., 2022, Taliaferro, Aspden et al., 2013, Tarallo, Giurato et al., 2017). In agreement with these canonical and non-canonical functions, AGO2 has been reported to shuttle between the cytosol and the nucleus of mammalian cells (Kalantari et al., 2016; Ohrt et al., 2008; Schraivogel and Meister, 2014). The relative subcellular distribution of AGO2 is also known to vary as a function of cell types and culture conditions, being notably generally increased in the nucleus of primary cell cultures as opposed to most immortalized cell lines (Sarshad et al., 2018; Sharma et al., 2016). Other conditions that increase AGO2 nuclear localization include cellular stress (Castanotto, Zhang et al., 2018, Rentschler, Chen et al., 2018), senescence (Benhamed et al., 2012; Rentschler et al., 2018) and differentiation (Sarshad et al., 2018). What is less clear is how AGO2 shuttles between both compartments. AGO2 does not contain a NLS or a nuclear export signal (NES). RISC-associated TNRC6 proteins (Trinucleotide repeat containing 6) contain both a NLS and a NES, thereby making them good candidates as intermediary factors. However, such a role appears limited to AGO2 export only, also involving the major nuclear export factor CRM1 (Chromosomal Maintenance 1) (Nishi, Nishi et al., 2013, Schraivogel, Schindler et al., 2015). As of today, the molecular mechanism of AGO2 nuclear import thus remains largely unknown. Although initial studies reported a role for Importin-8 (Weinmann, Hock et al., 2009), follow up work revealed that many importins are in fact redundantly involved (Schraivogel et al., 2015), and no NLS-containing protein is currently known to act as intermediary cargo for these importins.

Here, we report that the NLS-containing FAM172A regulates AGO2 nuclear import. Our data further suggest that perturbation of this process is clinically relevant, also impacting subsequent coregulation of alternative splicing by FAM172A and AGO2, independently of post-transcriptional gene silencing.

## Results

### FAM172A influences AGO2 nuclear localization

To determine if FAM172A plays a role in the nucleocytoplasmic shuttling of AGO2, we first compared the subcellular distribution of AGO2 in mouse embryonic fibroblasts (MEFs) freshly derived from *Fam172a*^*Tp/Tp*^ and wild-type (WT) e10.5 embryo heads – a cellular model that previously proved useful for highlighting the nuclear colocation of FAM172A and AGO2 proteins via immunofluorescence (Bélanger et al., 2018). As noted in this prior work, AGO2 is enriched in the nucleus of WT MEFs (Fig.1A-B), this enrichment being decreased when MEFs are plated at high density (Fig.S1A-B). Under both growth conditions (moderate and high density), we found that AGO2 is depleted from the nucleus of *Fam172a*^*Tp/Tp*^ MEFs thereby becoming markedly enriched in the cytoplasm (Figs.1A-B and S1A-B). A similar outcome was not seen for AGO1, which remains enriched in the nucleus of MEFs despite the loss of FAM172A (Fig.S1C-D). Using western blot and fractionation assays, we further confirmed that the loss of FAM172A specifically affects AGO2 nuclear localization and not overall protein levels (Figs.1C and S1E). Other immunofluorescence data revealed that the converse is not true: the predominantly nuclear localization of FAM172A is not altered by the presence of high levels of transfected _FLAG_AGO2 in the cytoplasm of control MEFs (Fig.S1F).

**Figure 1.**
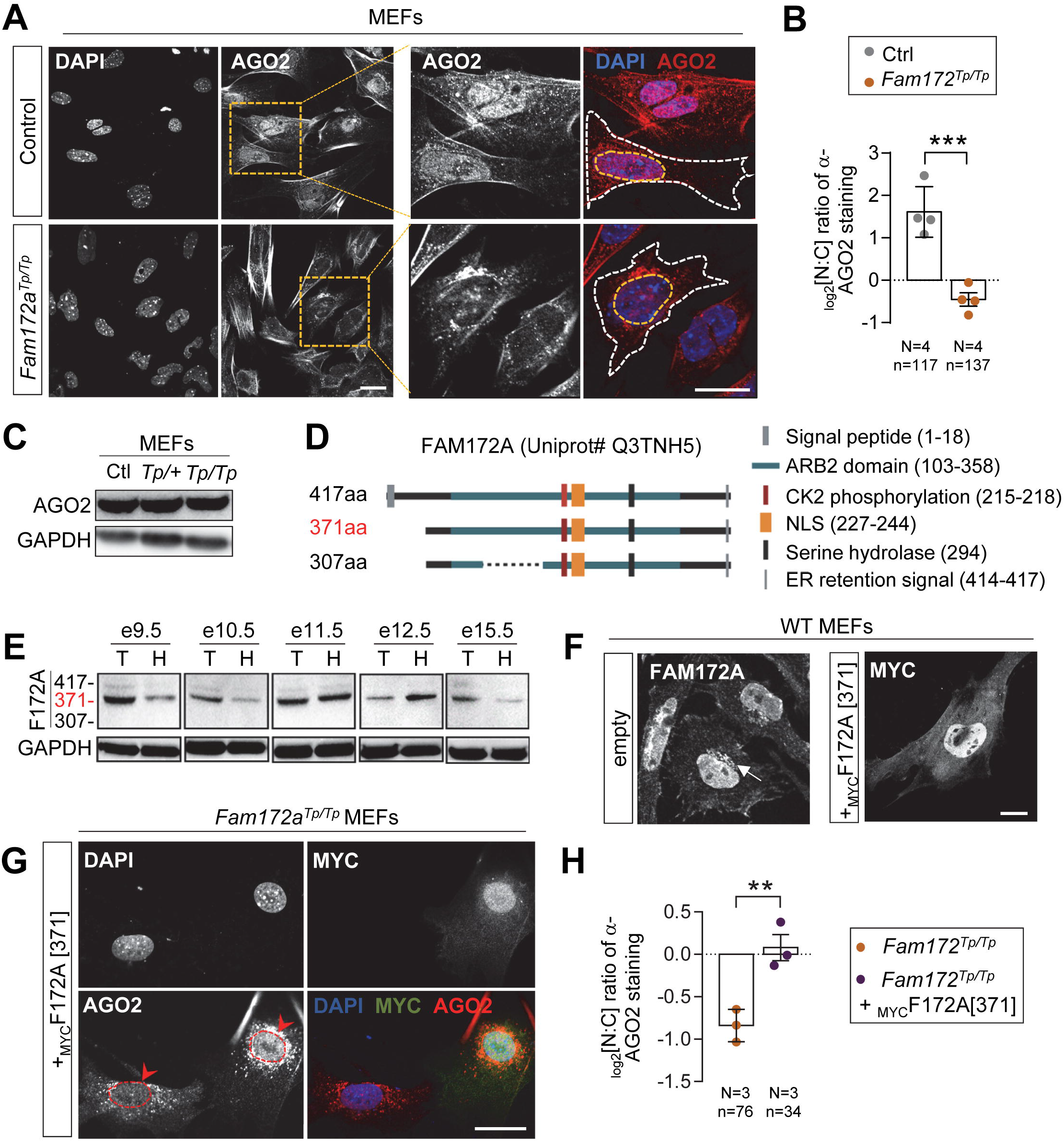
FAM172A influences AGO2 nuclear localization. **(A)** Immunofluorescence analysis of AGO2 distribution in WT and *Fam172*^*Tp/Tp*^ e10.5 MEFs at moderate density (40 000 cells/cm^2^), with nuclei stained using DAPI. **(B)** Quantification of relative fluorescence intensity of anti-AGO2 staining in nucleus and cytoplasm (N:C ratio, expressed in log 2 scale) using images such as those displayed in A. Yellow and white dashed lines in zoomed-in views in A delineate measured areas for nucleus and cytoplasm, respectively. **(C)** Western blot analysis of AGO2 protein levels in WT, *Fam172*^*Tp/+*^ and *Fam172*^*Tp/Tp*^ e10.5 MEFs, using GAPDH as loading control (N=3). **(D)** Diagram of mouse FAM172A isoforms based on Ensembl/Uniprot databases. **(E)** Western blot analysis of FAM172A isoforms in trunk (T) or head (H) extracts from embryos at indicated developmental stages, using GAPDH as loading control (N=3). **(F)** Immunofluorescence staining of endogenous FAM172A and transfected MYC-tagged version of the 371aa-long isoform (_**MYC**_F172A[371]) in e10.5 MEFs, with nuclei stained using DAPI (N=3). The arrow in the left panel points the endoplasmic reticulum. **(G)** Immunofluorescence analysis of AGO2 distribution in *Fam172*^*Tp/Tp*^ e10.5 MEFs transfected or not with _**MYC**_F172A[371]. Red arrowheads compare fluorescence intensity in nuclei of transfected and non-transfected cells. **(H)** Quantification of relative fluorescence intensity of anti-AGO2 staining in nucleus and cytoplasm (N:C ratio, expressed in log 2 scale) using images such as those displayed in G. Scale bar, 20μm. \*\**P* ≤ 0.01 and \*\*\**P* ≤ 0.001; Student’s *t*-test. (Further data can be found in Figure S1)

To strengthen our observations, we next sought to rescue AGO2 nuclear depletion in *Fam172a*^*Tp/Tp*^ MEFs via transfection of a FAM172A-expressing vector. To this end, we first reviewed the different isoforms of FAM172A, which have been updated since our previous work (Belanger et al., 2018). The Ensembl genome browser now predicts three major isoforms in mice (Fig.1D), all three being presumably detectable with the commercially-available polyclonal antibody used in our laboratory (Abcam ab121364). Using this antibody to probe embryo extracts at different developmental stages (e9.5 to e15.5), we found that the 371aa-long isoform is the most predominant, whereas the 417aa-and 307aa-long isoforms are weakly expressed and undetectable, respectively (Fig.1E). The same pattern was observed in several murine cell lines such as Neuro2a (N2a) neuroblasts, NIH 3T3 fibroblasts and R1 embryonic stem cells (Fig.S1G). Accordingly, we found that a MYC-tagged version of the 371aa isoform can recapitulate the subcellular distribution of endogenous FAM172A in MEFs (Fig.1F), except for the ER that is most likely targeted by the signal peptide-containing 417aa-long isoform (Bélanger et al., 2018). Upon transfection in *Fam172a*^*Tp/Tp*^ MEFs, this MYC-tagged version of the 371aa-long isoform (hereinafter referred to as _MYC_FAM172A) allowed to re-establish the predominantly nuclear localization of AGO2 (Fig.1G-H). Hence, we conclude that FAM172A is actively involved in the regulation of AGO2 nuclear localization.

### FAM172A directly interacts with AGO2

Co-immunoprecipitation (co-IP) assays in multiple cell types/tissues previously showed that FAM172A and AGO2 can be found in a same complex, but not when using a version of murine FAM172A mimicking the CHARGE syndrome-associated human variant pE228Q (pE229Q in mice) (Bélanger et al., 2018). To determine if the FAM172A-AGO2 interaction is direct, we performed *in vitro* co-IP assays using purified _MBP_FAM172A and _His_AGO2 recombinant proteins. Analysis of both WT and E229Q-mutated _MBP_FAM172A in this context revealed that the FAM172A-AGO2 interaction is direct and further confirmed the prominent role played by the NLS-located E229 residue of FAM172A (Fig.2A-B). To narrow down the interaction interface in AGO2, we also performed co-IP assays in N2a cells transfected with _MYC_FAM172A and deletion constructs of _FLAG_AGO2. This analysis identified the PAZ domain-containing N-terminal half of AGO2 as being essential for mediating the interaction with FAM172A (Fig.2C).

**Figure 2.**
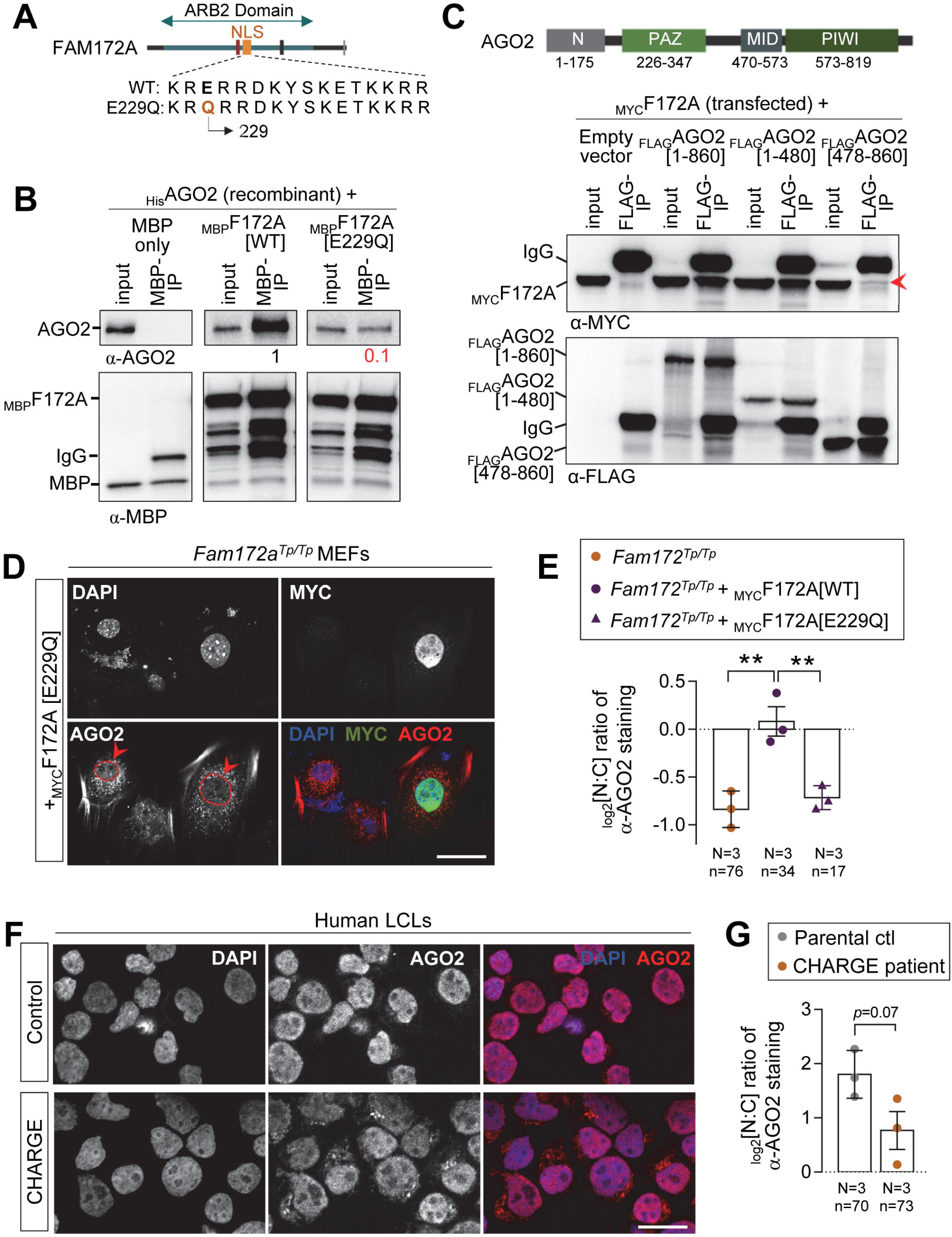
FAM172A directly interacts with AGO2. **(A)** Diagram of the E229Q point mutation in murine FAM172A protein sequence. **(B)** *In vitro* co-IP of recombinant _MBP_FAM172A (WT *vs* E229Q versions) and _His_AGO2 using MBP as bait (N=3). MBP alone was used as negative control. Indicated numbers correspond to relative amounts of co-immunoprecipitated _His_AGO2 after normalization for amount of immunoprecipitated _MBP_FAM172A, as determined via densitometry. **(C)** Co-IP of transfected _MYC_FAM172A and _FLAG_AGO2 (full-length *vs* indicated N-term and C-term truncations) in N2a cells using FLAG as bait (N=3). The red arrowhead points to reduced amount of co-immunoprecipitated _MYC_FAM172A when using the C-term half of AGO2. **(D)** Immunofluorescence analysis of AGO2 distribution in *Fam172*^*Tp/Tp*^ e10.5 MEFs transfected or not with E229Q-mutated _**MYC**_FAM172A, with nuclei stained using DAPI. Red arrowheads compare relative fluorescence intensity in nuclei of transfected and non-transfected cells. **(E)** Quantification of relative fluorescence intensity of anti-AGO2 staining in nucleus and cytoplasm (N:C ratio, expressed in log 2 scale) using images such as those displayed in D. Data for *Fam172*^*Tp/Tp*^ and *Fam172*^*Tp/Tp*^ + _MYC_F172A[WT] conditions are also shown in Figure 1H, being duplicated here for comparison purposes only. **(F)** Immunofluorescence analysis of AGO2 distribution in human LCLs derived from an individual with FAM172A[E228Q]-associated CHARGE syndrome and parental control, with nuclei stained using DAPI (N=3 technical replicates). **(G)** Quantification of relative fluorescence intensity of anti-AGO2 staining in nucleus and cytoplasm (N:C ratio, expressed in log 2 scale) using images such as those displayed in F. Scale bar, 20μm. \*\**P* ≤ 0.01; Student’s *t*-test. (Further data can be found in Figure S2)

To evaluate the functional relevance of this direct interaction for AGO2 nuclear localization, we then reiterated the rescue experiment in *Fam172a*^*Tp/Tp*^ MEFs. Strikingly, in contrast to WT _MYC_FAM172A (Fig.1G-H), we found that E229Q-mutated _MYC_FAM172A is unable to correct the abnormally enriched cytoplasmic distribution of AGO2 in these cells (Fig.2D-E). Of note, this lack of effect is not due to a reduced ability of E229Q-mutated _MYC_FAM172A to reach the nucleus (Fig.2D). Quantitative analyses in N2a cells even suggest that the E229Q mutation increases _MYC_FAM172A nuclear localization (Fig.S2A-B). Collectively, these data thus clearly show that AGO2 nuclear localization relies on a direct interaction between FAM172A and AGO2, which is dependent on the E229 residue of FAM172A and mediated by the PAZ domain-containing half of AGO2. Importantly, this mechanism is most likely at play in human cells as well, as evidenced by the reduced nuclear levels of AGO2 in a lymphoblastoid cell line (LCL) derived from a CHARGE syndrome child bearing the FAM172A variant E228Q (Fig.2F-G).

### The NLS of FAM172A is required for AGO2 nuclear import

To determine if FAM172A either plays an active role in AGO2 nuclear import or only prevents nuclear AGO2 to leave the nucleus, we turned to Bimolecular Fluorescence Complementation (BiFC) assays (Jia et al., 2021; Kerppola, 2008; Kodama and Hu, 2012). To this end, we generated a series of fusion proteins where the N-terminal half of Venus (a brighter variant of YFP; Yellow Fluorescent Protein) is located on either end of AGO2 and the C-terminal half of Venus is located on either end of FAM172A. Upon transfection in N2a cells, 3 out of 4 combinations allowed to reconstitute a fluorescent Venus protein (Figs.3A and S2C). The combination where both AGO2 and FAM172A have their respective Venus moiety at their N-terminal end (_N-Venus_AGO2 and _C-Venus_FAM172A) gave a more robust fluorescent signal and was thus selected for further studies (Fig.3A). Using flow cytometry, we confirmed the specificity of this BiFC signal using an E229Q-mutated version of _C-Venus_FAM172A as negative control (Fig.S2D) and by competition with an excess of _FLAG_AGO2 (Fig.S2E). Using immunofluorescence, we also verified that the non-fluorescent fragments fused to AGO2 and FAM172A do not alter their respective subcellular distribution in N2a cells (Fig.S2F). Initial observations of the subcellular distribution of the BiFC signal generated by the _N-Venus_AGO2–_C-Venus_FAM172A complex in living N2a and NIH 3T3 cells showed an enrichment in the cytoplasm after 48h of culture (Fig.S2C,G). This outcome was somehow surprising considering our previous co-IP experiments using N2a cell fractions (cytoplasm *vs* nucleus) that detected an AGO2-FAM172A complex in the nuclear fraction only (Bélanger et al., 2018). We interpret this apparent discrepancy to the methods used, with BiFC being able to capture and lock transient interactions that cannot be detected using co-IP (Jia et al., 2021; Kodama and Hu, 2012).

**Figure 3.**
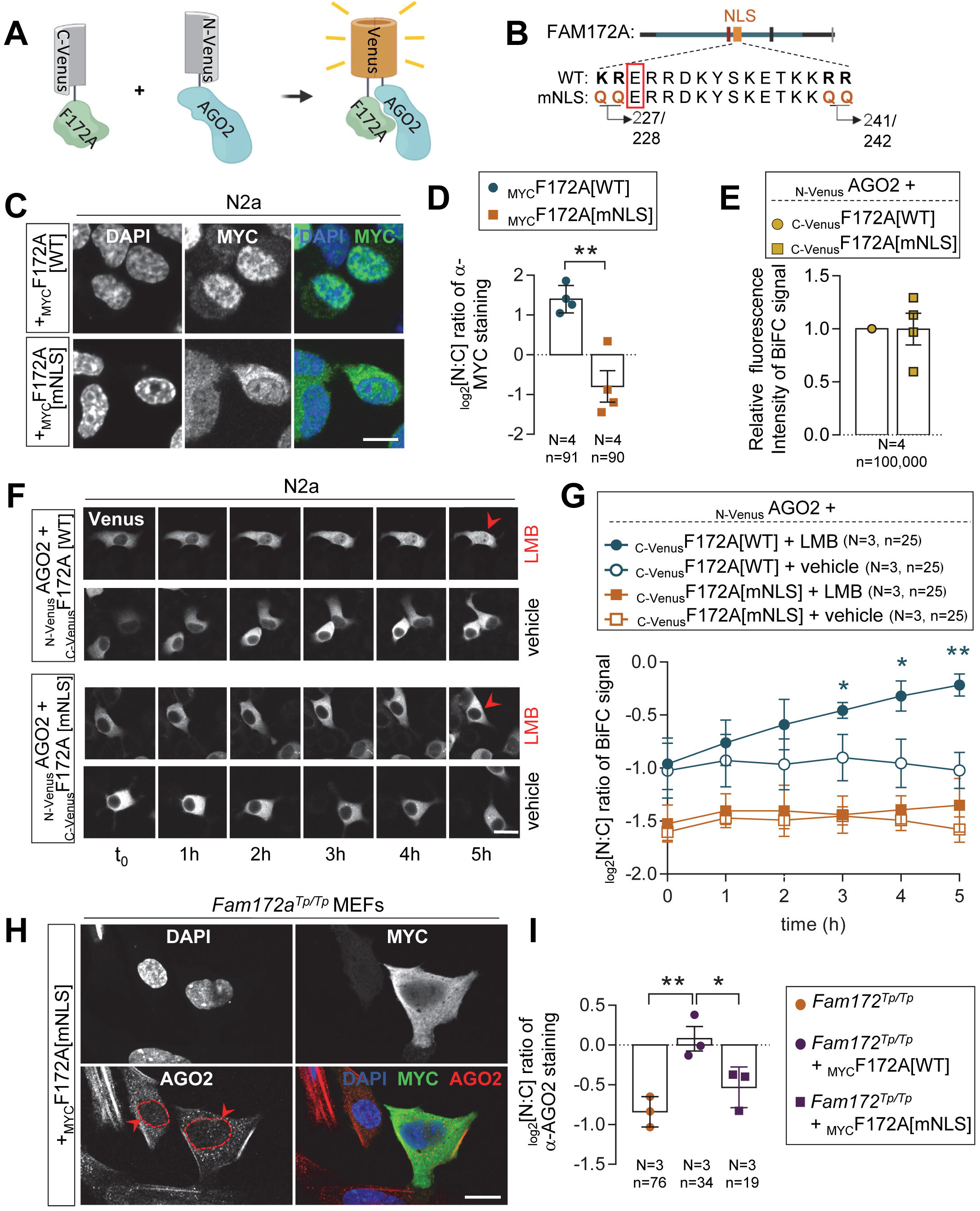
The NLS of FAM172A is required for AGO2 nuclear import. **(A)** Diagram of the BiFC assay based on structural complementation between two non-fluorescent N-terminal and C-terminal halves of a Yellow Fluorescent Protein (Venus) that are respectively fused to AGO2 (_N-Venus_AGO2) and FAM172A (_C-Venus_F172A). **(B)** Diagram of NLS point mutations (mNLS) in murine FAM172A protein sequence. The red box indicates the E229 residue mutated in some individuals with CHARGE syndrome. **(C)** Immunofluorescence analysis of MYC-tagged WT and NLS-mutated FAM172A proteins in transfected N2a cells, with nuclei stained using DAPI. **(D)** Quantification of relative fluorescence intensity of anti-MYC staining in nucleus and cytoplasm (N:C ratio, expressed in log 2 scale) using images such as those displayed in C. Data for _MYC_F172A[WT] condition are also shown in Figure S2B, being duplicated here for comparison purposes only. **(E)** Comparison of BiFC fluorescence intensity between _N-Venus_AGO2–_C-Venus_F172A[WT] and _N-Venus_AGO2–_C-Venus_F172A[mNLS]. Mean fluorescence intensity was determined using flow cytometry 24h after transfection in N2a cells. **(F)** Five-hour-long time-lapse recordings of BiFC signal generated using _N-Venus_AGO2-_C-Venus_F172A[WT] and _N-Venus_AGO2-_C-Venus_F172A[mNLS] in N2a cells treated with Leptomycin B (LMB) or vehicle (ethanol) only. Red arrowheads compare fluorescence intensity in nuclei of BiFC-positive cells. **(G)** Quantification of relative BiFC signal intensity in nucleus and cytoplasm (N:C ratio, expressed in log 2 scale) using images such as those displayed in F. **(H)** Immunofluorescence analysis of AGO2 distribution in *Fam172*^*Tp/Tp*^ e10.5 MEFs transfected or not with _**MYC**_F172A[mNLS], with nuclei stained using DAPI. Red arrowheads compare relative fluorescence intensity in nuclei of transfected and non-transfected cells. **(I)** Quantification of relative fluorescence intensity of anti-AGO2 staining in nucleus and cytoplasm (N:C ratio, expressed in log 2 scale) using images such as those displayed in H. Data for *Fam172*^*Tp/Tp*^ and *Fam172*^*Tp/Tp*^ + _MYC_F172A[WT] conditions are also shown in Figure 1H, being duplicated here for comparison purposes only. Scale bar, 10μm (C) and 20μm (F, H). \**P* ≤ 0.05 and \*\**P* ≤ 0.01; Student’s *t*-test.

The cytoplasmic enrichment of the BiFC signal supported the hypothesis that AGO2 and FAM172A initiate their interaction in the cytosol and then together enter the nucleus using the NLS of FAM172A. To test this model, we devised an experiment based on time-lapse imaging and blockage of CRM1-mediated nuclear export, also comparing WT and NLS-mutated versions of _C-Venus_FAM172A. In preparation for this experiment, we carefully point-mutated basic residues in each half of the bipartite NLS sequence of FAM172A (Fig.3B) in order to impair its nuclear import (Fig.3C-D) without affecting its capacity to interact with AGO2 (Fig.3E). N2a cells co-transfected with _N-Venus_AGO2 and either WT or NLS-mutated versions of _C-Venus_FAM172A were then followed under a confocal microscope during 5h after addition of the CRM1 inhibitor Leptomycin B (LMB). Remarkably, LMB addition led to a constant increase of the BiFC signal in the nucleus over time, but only when using WT _C-Venus_FAM172A (Fig.3F-G, Movie S1). With NLS-mutated _C-Venus_FAM172A, the BiFC signal already appeared more enriched in the cytoplasm before LMB addition and remained as such after LMB addition (Fig.3F-G, Movie S2). Altogether these data are consistent with the notion that AGO2 nuclear import is mediated by the NLS of its interacting partner FAM172A, as further demonstrated by the inability of an NLS-mutated version of _MYC_FAM172A to re-establish AGO2 nuclear enrichment in *Fam172a*^*Tp/Tp*^ MEFs (Fig.3H-I).

### AGO2 nuclear import depends on the status of FAM172A phosphorylation

The FAM172A interactome in N2a cells is known to include the catalytic subunit alpha of Casein kinase II (CK2) (Bélanger et al., 2018). Moreover, we noted that FAM172A bears two conserved consensus-like sequences for CK2 phosphorylation (S/T–X–X–D/E) close to its NLS (Fig.4A), and both publicly available (www.phosphosite.org) and our own (Fig.4B) mass spectrometry data confirm that the serine-rich motif (S215-S-S-S-D in mouse; S215-S-S-D in human) can indeed be phosphorylated. Given that proximity of a CK2 phosphorylation site was previously reported to influence NLS recognition by importins (Hübner et al., 1997; Jans et al., 2000; Xiao et al., 1997), we thus wondered if the status of CK2-mediated phosphorylation of FAM172A might play a regulatory role in the nuclear import of AGO2. To evaluate this possibility, we generated phosphodead (P-) and phosphomimetic (P+) versions of murine FAM172A, in which either the aspartate residues D220/D223 were replaced by glutamines (Litchfield, 2003, Lussier, Gu et al., 2014) or the upstream serine/threonine residues were replaced by aspartates (Sun, Cui et al., 2016), respectively (Fig.4A). These new constructs were then tested for their ability to mediate AGO2 nuclear import using the BiFC system in living N2a cells. With the phosphodead _C-Venus_FAM172A, we found that the BiFC signal generated in conjunction with _N-Venus_AGO2 remains enriched in the cytoplasm regardless of the presence of LMB or not (Fig.4C-D, Movie S3) – just like when we used the NLS-mutated _C-Venus_FAM172A (Fig.3F-G, Movie S2). In stark contrast, the BiFC signal appears greatly enriched in the nucleus when using the phosphomimetic _C-Venus_FAM172A (Fig.4C-D, Movie S4), with both basal and LMB-induced levels being greater than for WT _C-Venus_FAM172A (Figs.3F-G and 4D, Movie S2). We further observed that both phosphodead and phosphomimetic versions of _C-Venus_FAM172A do not affect overall intensity of the _N-Venus_AGO2–_C-Venus_FAM172A BiFC signal, only its cytoplasmic-*vs*-nuclear distribution (Fig.4E). Yet, analysis of the subcellular distribution of phosphodead and phosphomimetic versions of _MYC_FAM172A also showed that CK2-mediated phosphorylation of FAM172A is stimulatory but apparently not essential for its own entry in the nucleus (Fig.S3A-B). Nonetheless, this series of experiments strongly suggests that CK2-mediated phosphorylation of FAM172A is regulatory important for FAM172A-dependent nuclear import of AGO2.

**Figure 4.**
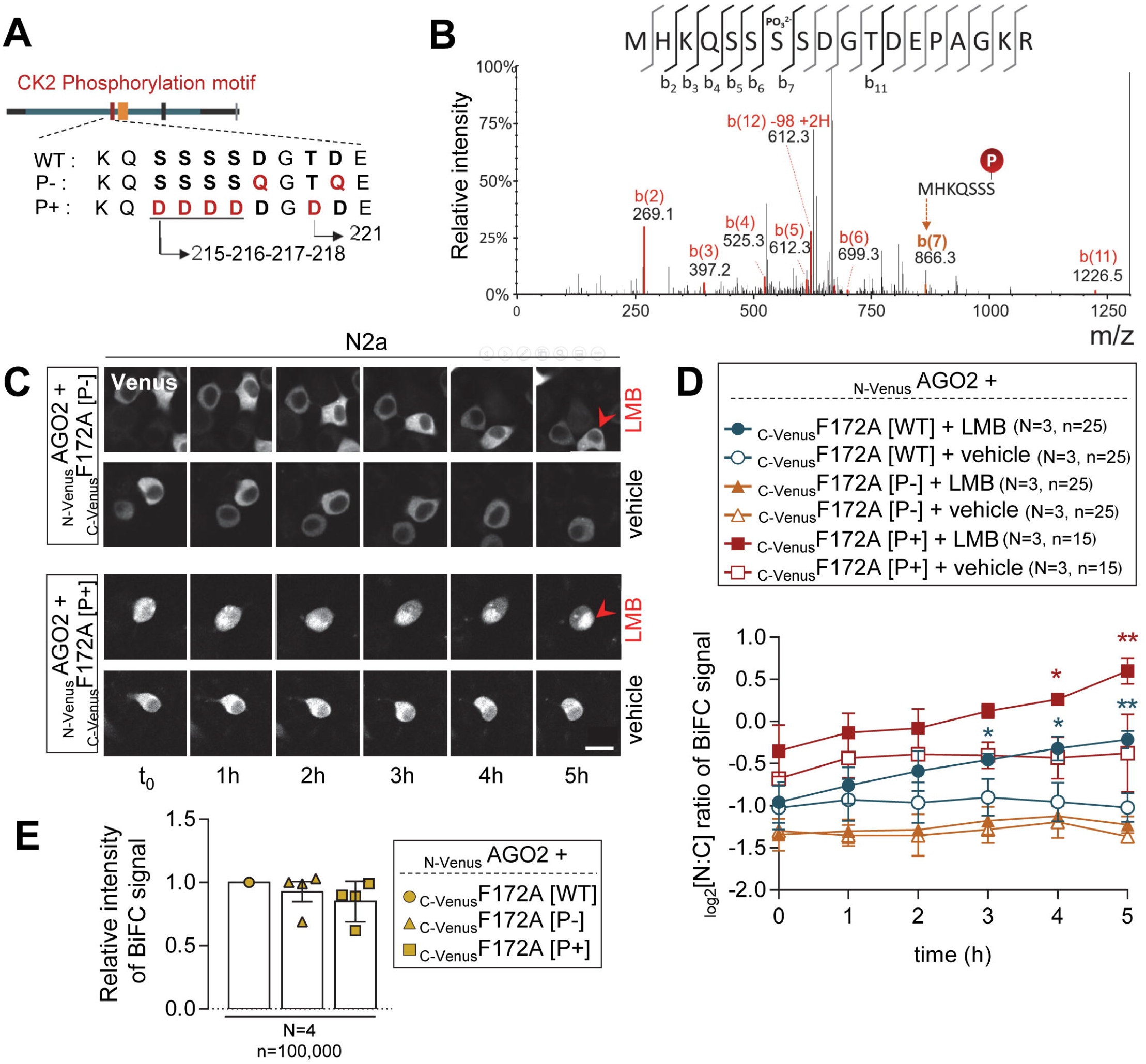
AGO2 nuclear import depends on the status of FAM172A phosphorylation. **(A)** Diagram of phosphodead (P[-]) and phosphomimic (P[+]) mutations in murine FAM172A protein sequence. **(B)** Representative CID-MS/MS spectrum of a phosphorylated FAM172A peptide on position S217, recovered after anti-MYC immunoprecipitation from _MYC_FAM172A-transfected N2a cells. **(C)** Five-hour-long time-lapse recordings of BiFC signal generated using _N-Venus_AGO2– _C-Venus_F172A[P-] and _N-Venus_AGO2–_C-Venus_F172A[P+] in N2a cells treated with Leptomycin B (LMB) or vehicle (ethanol) only. Red arrowheads compare fluorescence intensity in nuclei of BiFC-positive cells. **(D)** Quantification of relative BiFC signal intensity in nucleus and cytoplasm (N:C ratio, expressed in log 2 scale) using images such as those displayed in C. Data for _C-Venus_F172A[WT] are also shown in Figure 3G, being duplicated here for comparison purposes only. **(E)** Comparison of overall BiFC fluorescence intensity between _N-Venus_AGO2–_C-Venus_F172A[WT], _N-Venus_AGO2–_C-Venus_F172A[P-] and _N-Venus_AGO2–_C-Venus_F172A[P+]. Mean fluorescence intensity was determined using flow cytometry 24h after transfection in N2a cells. Scale bar, 20μm (C). \**P* ≤ 0.05 and \*\**P* ≤ 0.01; Student’s *t*-test. (Further data can be found in Figure S3)

### AGO2, together with FAM172A, fulfills a non-canonical function relevant for CHARGE syndrome pathogenesis

Data from our prior work (Bélanger et al., 2018) and the current study strongly suggested a role for AGO2 misregulation in CHARGE syndrome pathogenesis. To directly evaluate this possibility, we generated conditional mutants using a floxed allele of *Ago2* (O’Carroll, Mecklenbrauker et al., 2007) and the *Wnt1-Cre2* driver (Lewis, Vasudevan et al., 2013) – thereby allowing to test AGO2 function in the cell lineage most relevant for CHARGE syndrome (*i*.*e*., NCCs) while also circumventing the early embryonic lethality associated with constitutive loss of *Ago2* (Liu, Carmell et al., 2004, Morita, Horii et al., 2007). Resulting homozygous mutants (*Wnt1-Cre2*^*Tg/+*^*;Ago2*^*Flox/Flox*^; hereinafter referred to as *Ago2*^*cKO/cKO*^) were unable to breathe properly and therefore died at birth (Figs.5A-B and S4A, Movie S5), most likely because of fully penetrant bilateral choanal atresia (Fig.5C) – a cardinal feature of CHARGE syndrome. Other potential causes of respiratory failure that might have involved abnormal lung/diaphragm morphology and/or innervation were not found in these mutants (Fig.S4B-D). In addition to choanal atresia, *Ago2*^*cKO/cKO*^ embryos displayed different combinations of supplemental CHARGE syndrome-associated anomalies (Table S1), including coloboma (Fig.5D-E), retarded growth (Fig.5F), dysmorphism of external ears (Fig.S4E) and cranial nerves (Fig.S4F), and thymus hypoplasia (Fig.S4G-H). Of note, coloboma was found to mainly affect one eye only, with a small subset of animals also displaying microphthalmia (Fig.5D). These results convincingly show that the loss of *Ago2* in NCCs phenocopies CHARGE syndrome in mice, with choanal atresia, coloboma and thymus hypoplasia being the most highly penetrant features.

**Figure 5.**
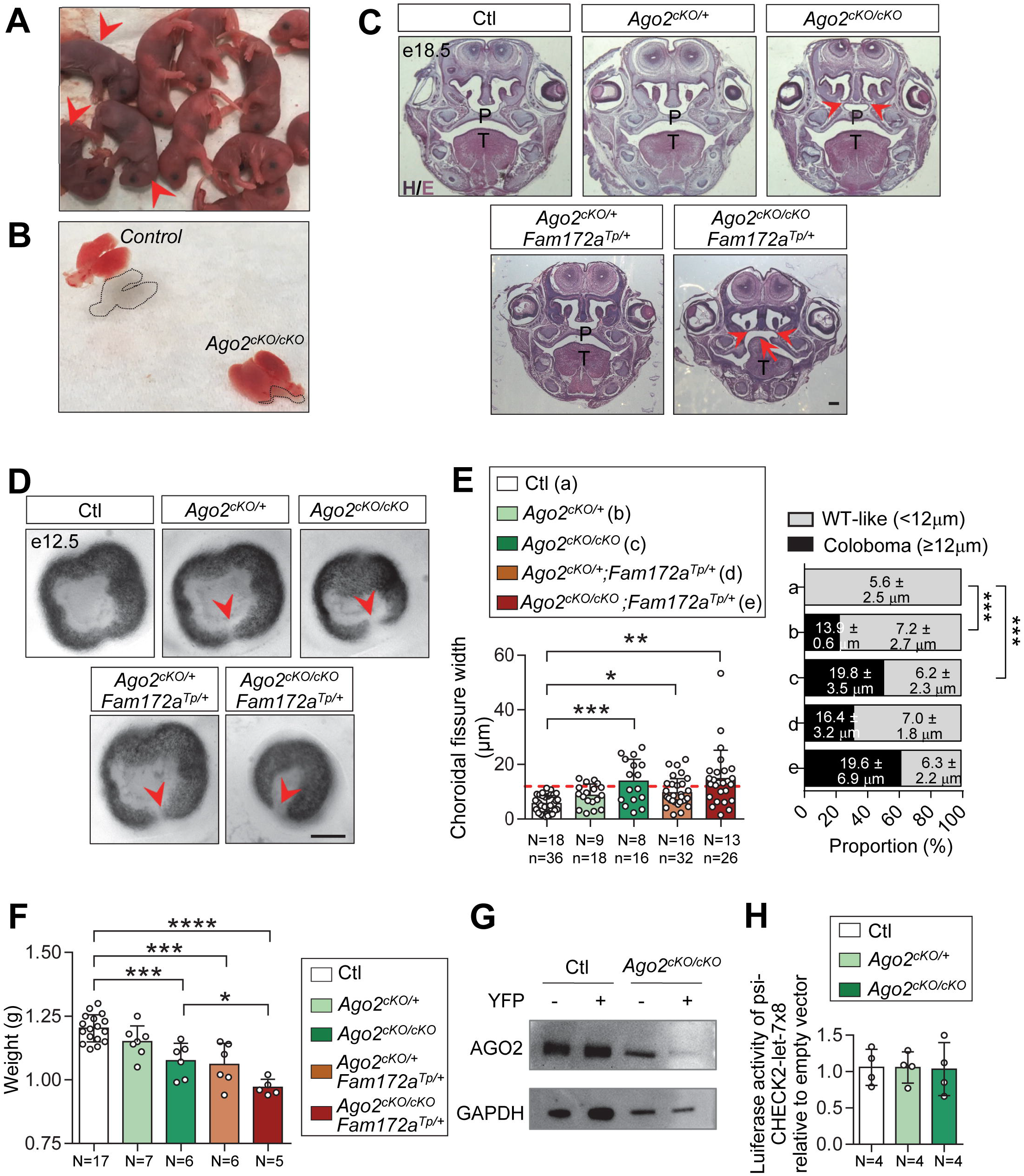
AGO2, together with FAM172A, fulfills a non-canonical function relevant for CHARGE syndrome pathogenesis. **(A-B)** *Ago2*^*cKO/cKO*^ newborns are cyanotic (red arrowheads in A) and lack air in their lungs (see floating test in B, with shadow of floating control lung delineated by dashed outline). **(C)** H&E-stained cross-sections of e18.5 fetus heads of indicated genotypes, with red arrowheads indicating blockage of nasal passages (choanal atresia) and red arrow indicating cleft palate (N=5 per genotype). P,palate, T, tongue. **(D)** Bright field view of e12.5 eyes of indicated genotypes, with red arrowheads pointing to unclosed choroidal fissure (coloboma). **(E)** Quantitative analysis of fissure width (left panel) and percentage of coloboma occurrence (right panel), using images such as those displayed in D. Red dotted line in left panel corresponds to the minimal width for coloboma (12μm; based on the maximum width observed in control embryos). The average width of choroidal fissure for coloboma and WT-like subgroups are indicated in their corresponding bar subdivisions in right panel. **(F)** Body weight of e18.5 fetuses of indicated genotype. **(G)** Western blot analysis of AGO2 protein levels in FACS-recovered cells (both YFP-negative and -positive) from e10.5 control (*Wnt1-Cre2*^*Tg/+*^;*Rosa26*^*FloxSTOP-YFP/+*^) and homozygous *Ago2* mutant (*Wnt1-Cre2*^*Tg/+*^;*Rosa26*^*FloxSTOP-YFP/+*^*;Ago2*^*Flox/Flox*^) embryos, with GAPDH used as loading control. **(H)** Luciferase assay in YFP-positive cells recovered by FACS from e10.5 embryos (owing to the *Rosa26*^*FloxSTOP-YFP*^ reporter) and transfected with either the psi-CHECK2-let-7×8 reporter or empty psi-CHECK2 vector. Scale bar, 1 mm (B) and 50 μm (C). \**P* ≤ 0.05, \*\**P* ≤ 0.01, \*\*\**P* ≤ 0.001 and \*\*\*\**P* ≤ 0.0001; Khi2 test (E, right) or one-way ANOVA with Tukey’s post-hoc test. (Further data can be found in Table S1 and Figure S4)

To strengthen the link with CHARGE syndrome, we then tested a genetic interaction with *Fam172a*. Strikingly, we found that addition of a single *Toupee* allele of *Fam172a* – which is not sufficient by itself to cause a phenotype (Bélanger et al., 2018) – was here sufficient to worsen the phenotypes of both heterozygous and homozygous *Ago2* conditional mutants (Table S1). Notably, all *Ago2*^*cKO/cKO*^*;Fam172a*^*Tp/+*^ fetuses were found to display cleft palate in addition to choanal atresia (Fig.5C). Head of these fetuses also appeared more flat compared to single *Ago2*^*cKO/cKO*^ mutants (Fig.5C). Moreover, we observed a clear gene dosage-dependent impact on the frequency and severity of growth retardation (Fig.5F), while the impact of the *Fam172a*^*Tp*^ allele on ocular defects was best evidenced by a robust increase in the frequency of bilateral coloboma (from 14% to 89%) and association with microphthalmia (from 14% to 44%) in *Ago2*^*cKO/cKO*^*;Fam172a*^*Tp/+*^ embryos compared to *Ago2*^*cKO/cKO*^ embryos (Fig.5D and Table S1). As *Fam172a* expression is not affected by the loss of AGO2 (Fig. S4I), these *Ago2-Fam172a* genetic interaction data are in agreement with a shared role for AGO2 and FAM172A in NCCs that is relevant for CHARGE syndrome pathogenesis.

In parallel studies, we also generated *Ago2*^*cKO/cKO*^ embryos bearing the *Rosa26*^*FloxSTOP-YFP*^ reporter allele in order to confirm the NCC-specific loss of AGO2. This allowed us to recover YFP-positive NCCs and YFP-negative non-NCCs by FACS (fluorescence-activated cell sorting) from e10.5 embryos, which were subsequently analyzed using western blot. As expected, this analysis revealed a robust and specific decrease of AGO2 protein levels in NCCs from *Ago2*^*cKO/cKO*^ mutant embryos compared to controls (Fig.5G). Most importantly, we further took advantage of this system to verify the status of the canonical AGO-dependent miRNA effector pathway in e10.5 NCCs. As previously observed in *Fam172a*^*Tp/Tp*^ MEFs (Belanger et al., 2018), transfection of the psi-CHECK2-let-7 × 8 luciferase reporter (Johnston, Geoffroy et al., 2010) in these FACS-recovered cells showed that AGO2 is dispensable for cytoplasmic post-transcriptional gene silencing in this context (Fig.5H) – most likely because of functional redundancy with other AGO proteins (Su, Trombly et al., 2009). From all these studies in mice, we thus conclude that AGO2-FAM172A functional interplay is needed in the nucleus of NCCs to prevent the development of CHARGE syndrome-associated anomalies, independently of canonical miRNA signaling.

### AGO2 and FAM172A coregulate alternative splicing *in vivo*

One key non-canonical function of AGO2 in the nucleus is to regulate cotranscriptional alternative splicing (Ameyar-Zazoua et al., 2012, Batsche & Ameyar-Zazoua, 2015, Chu et al., 2021, Kalantari et al., 2016, Taliaferro et al., 2013, Tarallo et al., 2017), which we previously showed to be dysregulated in the context of CHARGE syndrome in both mouse models and patient-derived LCLs (Belanger et al., 2018). To verify if a similar mechanism could explain the CHARGE syndrome-associated anomalies in *Ago2*^*cKO/cKO*^ embryos (Figs.5 and S4; Table S1), we analyzed a set of known splicing targets of AGO2 (Ameyar-Zazoua et al., 2012) that were also found to be dysregulated by the loss of either FAM172A or CHD7 (Belanger et al., 2018). RT-qPCR analysis revealed that virtually all of the eight tested splicing events for this set of four genes (*Mical2, Ift74, Col5a3*, and *Cd44*) were dysregulated in e12.5 *Ago2*^*cKO/cKO*^ heads (Fig.6A), with only two splicing events missing the statistical cut-off by little. To consolidate our hypothesis of a collaborative function for AGO2 and FAM172A, we also analyzed the same set of splicing targets in *Ago2-Fam172* double-mutants. In line with the observed gene dosage-dependent impact on the severity of CHARGE syndrome-associated anomalies (Fig.5C-F; Table S1), presence of a single *Fam172a*^*Tp*^ allele appeared generally sufficient to worsen the abnormal splicing signature associated with either heterozygous or homozygous loss of AGO2 (Fig.6A).

**Figure 6.**
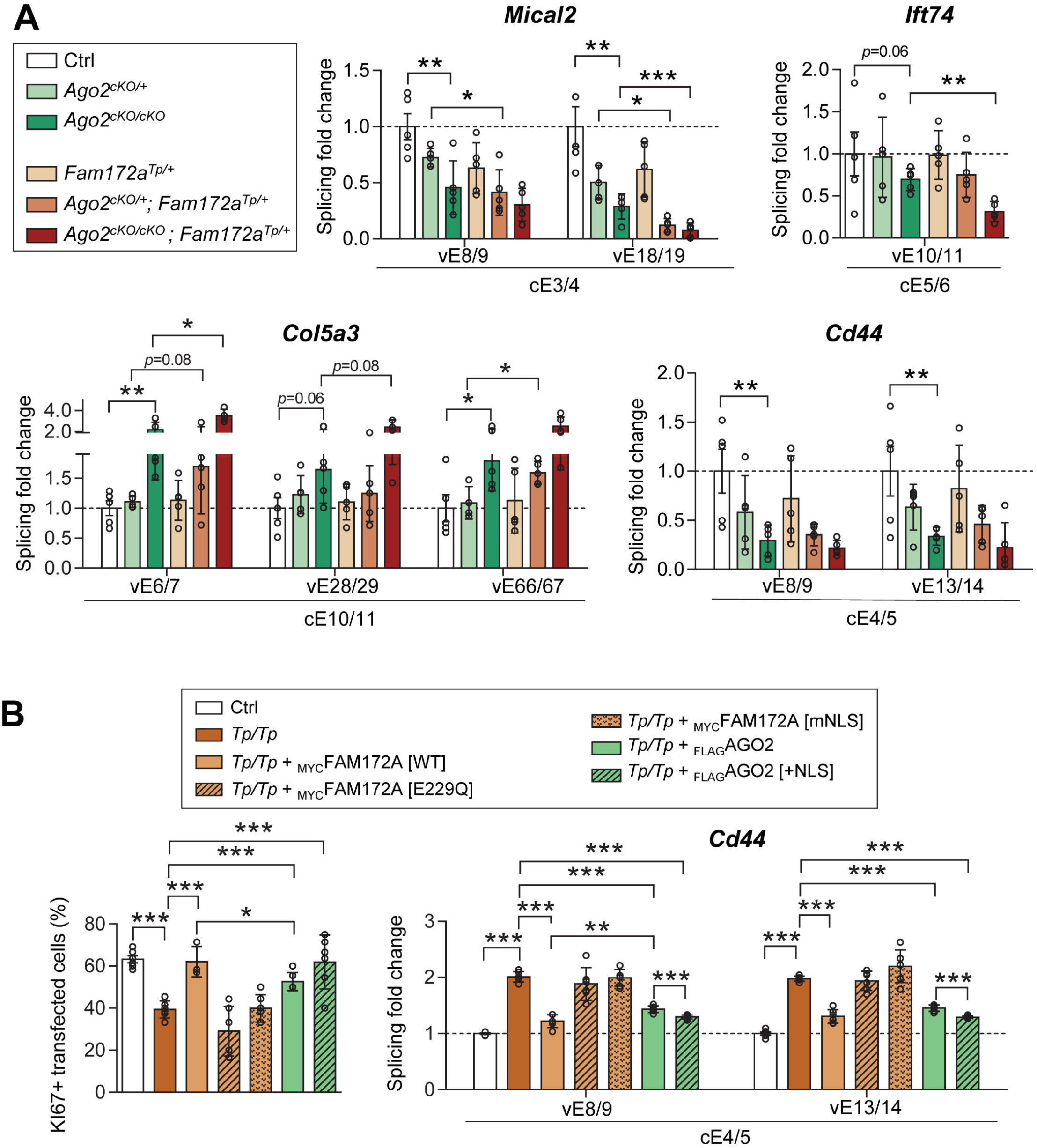
AGO2 and FAM172A coregulate alternative splicing *in vivo*. **(A)** RT-qPCR analysis of alternative splicing events for *Mical2, Ift74, Col5a3* and *Cd44* in e12.5 heads from an allelic series of *Ago2*^*cKO*^ and *Fam172a*^*Tp*^ mutant embryos. Expression levels of indicated variable exon pairs (vE) were normalized against levels of a constant exon pair (cE), and splicing fold change was then determined by comparison to the reference value (dashed line) obtained with WT control embryos (N=5 per genotype). **(B)** Quantification of cell proliferation (left panel) and alternative splicing of *Cd44* (right panel) in e10.5 *Fam172a*^*Tp/Tp*^ MEFs after cotransfection with a GFP expression vector and either _MYC_FAM172A (WT, E229Q-mutant or NLS-mutant) or _FLAG_AGO2 (WT or NLS-containing) expression vectors. Cell proliferation is expressed in % of GFP-positive cells based on double-immunofluorescence staining of GFP and the proliferation marker Ki67. Alternative splicing of *Cd44* was analyzed as described above, after FACS-mediated recovery of GFP-positive cells. \**P* ≤ 0.05, \*\**P* ≤ 0.01, \*\*\**P* ≤ 0.001; Student’s *t*-test.

In addition, we further tested the potential collaborative function of AGO2 and FAM172A via rescue experiments in freshly dissociated *Fam172a*^*Tp/Tp*^ MEFS, using proliferation rate and alternative splicing of *Cd44* as functional readouts (Fig.6B). To this end, *Fam172a*^*Tp/Tp*^ MEFS were transfected with either _MYC_FAM172A (WT, E229Q-mutant or NLS-mutant) or _FLAG_AGO2 (WT or NLS-containing), and then recovered by FACS owing to coexpressed GFP. As previously observed (Belanger et al., 2018), transfection of WT _MYC_FAM172A fully rescued the otherwise decreased proportion of proliferating (Ki67-positive) *Fam172a*^*Tp/Tp*^ MEFS compared to WT MEFs (Fig.6B, left). In contrast, both E229Q-mutated and NLS-mutated versions were totally unable to do so (Fig.6B, left), consistent with their incapacity to rescue AGO2 nuclear import in these cells; Figs.2D-E and 3H-I). Interestingly, overexpression of WT _FLAG_AGO2 (which increases overall levels in both cytoplasm and nucleus; Fig.S1F) partially rescued while another version engineered to contain a strong NLS (3 copies of SV40 LT monopartite NLS; Fig.S5A) fully rescued the proliferation rate of *Fam172a*^*Tp/Tp*^ MEFS (Fig.6B, left). Similar results were obtained when assessing *Cd44* alternative splicing (Fig.6B, right). Altogether, these data are thus in agreement with a general model suggesting that proper FAM172A-mediated nuclear import of AGO2 and subsequent coregulation of alternative splicing by FAM172A and AGO2 are necessary to avoid developing CHARGE syndrome.

### Acute rapamycin treatment rescues alternative splicing defects and coloboma in *Ago2*- and *Fam172a*-mutant embryos

Owing to its potent capacity to downregulate TOR-dependent ribosomal protein gene expression, rapamycin can promote splicing of other pre-mRNAs by relieving competition for a limiting pool of splicing factors (Munding, Shiue et al., 2013). Based on this premise, we previously used acute rapamycin treatment to correct alternative splicing defects in CHARGE syndrome patient-derived LCLs and coloboma in *Fam172a*^*Tp/Tp*^ mouse embryos (Belanger et al., 2018). Yet, impact on alternative splicing was not formally determined in these embryos. To address this lack and also determine the contribution of dysregulated alternative splicing in the development of CHARGE syndrome-related anomalies in *Ago2*^*cKO/cKO*^ embryos, we thus revisited our rapamycin treatment approach.

Initial experiments performed using the same treatment regimen as before (i.p. injection of 1mg/kg rapamycin to pregnant dams, twice a day during the peak of NCC migration between e9.5 to 11.5) severely affected general growth of *Ago2*^*cKO/cKO*^ embryos (Fig.S5B). This growth delay at e12.5 appeared more severe than previously observed for *Fam172a*^*Tp/Tp*^ embryos (Belanger et al., 2018), most likely because AGO2 loss and associated anomalies are restricted to a single cell lineage – thereby making all other cell types vulnerable to the teratogenic effects of rapamycin in a WT context (Hentges, Sirry et al., 2001). Decreasing rapamycin dose and frequency to 0.5 mg/kg once daily during the same period of time allowed us to limit these adverse effects at e12.5 (Fig.S5C). Importantly, this low-dose rapamycin treatment also improved the closure of the choroidal fissure, thereby markedly reducing coloboma incidence in *Ago2*^*cKO/cKO*^ embryos (Fig.7A-B) – in a way similar to what we previously observed for *Fam172a*^*Tp/Tp*^ embryos treated with the initial higher but still moderate dose (Belanger et al., 2018). Accordingly, most of the alternative splicing events for *Mical2, Ift74, Col5a3*, and *Cd44* that were tested at e12.5 were found to be similarly corrected (at least partially) in both mutants (Fig.7C). These *in vivo* data further lend credence to the previously proposed causal role of global dysregulation of alternative splicing in CHARGE syndrome pathogenesis (Belanger et al., 2018, Berube-Simard & Pilon, 2018).

**Figure 7.**
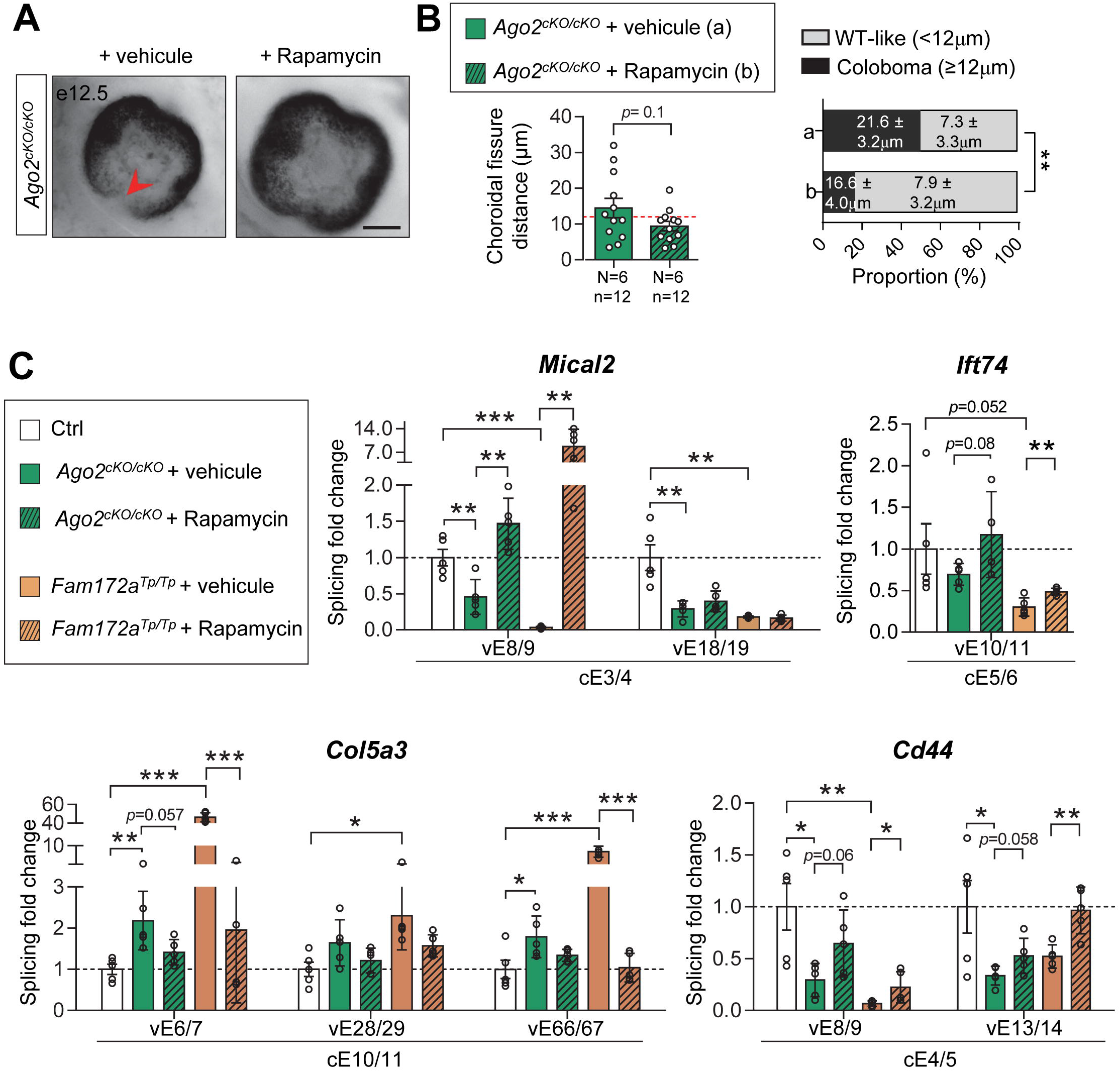
Acute rapamycin treatment rescues alternative splicing defects and coloboma in *Ago2*- and *Fam172a*-mutant embryos. **(A)** Bright field views of e12.5 eyes from *Ago2*^*cKO/cKO*^ embryos that were exposed to either rapamycin (0.5 mg/kg body weight) or vehicle (20% ethanol) *in utero* between e9.5 to e11.5. The red arrowhead indicates an unclosed choroidal fissure (coloboma). **(B)** Quantitative analysis of fissure width (left panel) and percentage of coloboma occurrence (right panel), using images such as those displayed in A. The red dotted line in left panel corresponds to the minimal width for coloboma (12μm; based on the maximum width observed in control embryos). The average width of choroidal fissure for coloboma and WT-like subgroups are indicated in their corresponding bar subdivisions in right panel. **(C)** RT-qPCR analysis of splicing events for *Mical2, Ift74, Col5a3* and *Cd44* in e12.5 heads from *Ago2*^*cKO/cKO*^ and *Fam172a*^*Tp/Tp*^ embryos that were exposed to either rapamycin (0.5 mg/kg and 1mg/kg body weight for *Ago2*^*cKO/cKO*^ and *Fam172a*^*Tp/Tp*^, respectively) or vehicle (20% ethanol) *in utero* between e9.5 to e11.5. Expression levels of indicated variable exon pairs (vE) were normalized against levels of a constant exon pair (cE), and splicing fold change was then determined by comparison to the reference value (dashed line) obtained with WT control embryos (N=5 per condition). Scale bar, 50 μm. \**P* ≤ 0.05, \*\**P* ≤ 0.01 and \*\*\**P* ≤ 0.001; Khi2 test (B) or Student’s *t*-test (C). (Further data can be found in Figure S5)

## Discussion

To follow up on our prior work and learn more about the poorly characterized CHARGE syndrome-associated protein FAM172A, we here focused on its interaction with the small RNA-binding protein AGO2 – which is known to play key gene regulatory roles in the nucleus beyond its canonical function in post-transcriptional gene silencing in the cytoplasm (Nazer et al., 2022). Previously published models predicted a central role for AGO2 at the chromatin-spliceosome interface (Ameyar-Zazoua et al., 2012), together with FAM172A that appears required for stabilizing protein-protein interactions (Belanger et al., 2018, Berube-Simard & Pilon, 2018). Our new data generated using various complementary approaches in murine and patient-derived cells now suggest that FAM172A is in addition a critical regulator of AGO2 nuclear import, involving both the nuclear localization signal and CK2 phosphorylation motif of FAM172A. In agreement with these key roles, we further found that *Ago2* genetically interacts with *Fam172a* in mice and that NCC-specific loss of AGO2 phenocopies CHARGE syndrome via dysregulation of alternative splicing. Overall, this study thus unveiled a new aspect of AGO2 function, with clinical relevance not only for CHARGE syndrome but also potentially other related developmental syndromes.

### FAM172A is a key regulator of AGO2 nuclear import

Although AGO2 has long been known to shuttle between the cytosol and nucleus of mammalian cells (Kalantari et al., 2016, Ohrt, Mutze et al., 2008, Rudel, Flatley et al., 2008, Schraivogel & Meister, 2014), we still know very little about the underlying mechanisms. Previous studies revealed that many different NLS-binding importins are likely mediating AGO2 nuclear entry (Schraivogel et al., 2015, Weinmann et al., 2009), but no NLS-containing proteins were previously identified as intermediary factors for the NLS-free AGO2. Such a role has been tested for members of the well-known AGO2-binding protein family TNRC6, which contain both a NLS and a NES (Nishi et al., 2013), with results being consistent with a role in AGO2 nuclear export but not import (Nishi et al., 2013, Schraivogel et al., 2015). Notably, knockdown of importin-β was shown to specifically impair the nuclear import of TNRC6 proteins without perturbing AGO2 nuclear import (Schraivogel et al., 2015). In contrast, we now present several lines of evidence demonstrating a key role for FAM172A in this process, with BiFC and rescue data clearly involving its NLS (Fig.3). Importantly, in total agreement with the existence of multiple nuclear import routes for AGO2 (Schraivogel et al., 2015), an exploratory BioID2-based screen for FAM172A interactors (to be published elsewhere) also identified several different importins (*e*.*g*., IMP-α, -4, -5, -7, -9, -11) and nucleoporins (*e*.*g*., NUP-50, -88, -93, -98, -155, -205, -214). Our analyses of the E229Q-mutated version of murine FAM172A further suggests that the interaction interface for AGO2 partly overlaps the NLS of FAM172A. Yet, the AlphaFold-predicted 3D structure of FAM172A (with its NLS protruding away from the core; Fig.S6A) indicates that this region is nonetheless well able to accommodate interactions with both AGO2 and importins.

More intriguing is the fact that the loss of FAM172A in MEFs decreases nuclear levels of AGO2 (Fig.1A) but not those of the closely related AGO1 (Fig.S1C). These observations are consistent with our previous co-IP experiments where we failed to detect an interaction between transfected _MYC_FAM172A and endogenous AGO1 in COS7 cells, despite having used exact same conditions that worked for endogenous AGO2 (Belanger et al., 2018). However, new data show that FAM172A can interact with AGO1 under certain conditions, like when using recombinant proteins *in vitro* (Fig.S6B) or upon overexpression in N2a cells (Fig.S6C). A plausible explanation for all these observations might be that FAM172A simply has a greater affinity for AGO2 than for AGO1, as suggested by the decreased ability of _FLAG_AGO1 to co-immunoprecipitate _MYC_FAM172A in transfected N2A cells compared to _FLAG_AGO2 (Fig.S6C).

Interestingly, our data suggesting a regulatory role for CK2 in FAM172A-mediated AGO2 nuclear import (Fig.4) provide a potential mechanistic link to explain why and how different cellular stresses can increase AGO2 translocation toward the nucleus (Castanotto et al., 2018, Rentschler et al., 2018). Indeed, CK2 activity is known to be enhanced by multiple cellular stressors (Ampofo, Sokolowsky et al., 2013, Sayed, Kim et al., 2000, Watabe & Nakaki, 2011). Moreover, it is noteworthy that about half of the proteins (43%; 29/67 proteins) composing the stress-induced response complex (SIRC) associated with AGO2 in HEK293 cells (Castanotto et al., 2018) are also included in the published FAM172A interactome from N2a (Belanger et al., 2018) and/or PGR9E11 cells (Belanger et al., 2022). One protein identified as being of special significance in the SIRC is Nucleolin (Castanotto et al., 2018), which we further validated as interactor for both FAM172A and AGO2 in N2a cells using co-IP (Fig.S6D). All these observations further make sense when considering the particular vulnerability of NCCs to cellular stress in general, along with the extensive overlap between neurocristopathies and spliceosomopathies (Beauchamp, Alam et al., 2020, Berube-Simard & Pilon, 2018, Pilon, 2021). Stress-induced and FAM172A-mediated nuclear import of AGO2 might be a way for this unique, multipotent and highly migratory cell population to defend against cellular stressors.

### A non-canonical but nonetheless clinically relevant role for nuclear AGO2 in NCCs

Constitutive homozygous loss of AGO2 is incompatible with life in both humans and mice (Lessel, Zeitler et al., 2020, Morita et al., 2007), most likely due to a combination of defective gastrulation (Alisch, Jin et al., 2007) and impaired differentiation of the extra-embryoni endoderm necessary for proper yolk sac formation (Ngondo, Cirera-Salinas et al., 2018). These critical functions are not shared with any other AGO family members (Modzelewski, Holmes et al., 2012, Van Stry, Oguin et al., 2012), and this is apparently not due to the unique endonuclease activity of AGO2 (Cheloufi, Dos Santos et al., 2010, Liu et al., 2004, Ngondo et al., 2018). A Cre/LoxP-based approach allowed us to circumvent this embryonic lethality in mice and directly test a potential role for AGO2 in CHARGE syndrome pathogenesis. In total accordance with this possibility, NCC-specific homozygous loss of AGO2 resulted in multiple cardinal features of CHARGE syndrome, including choanal atresia, coloboma and growth retardation (Fig.5). Of note, a role for AGO2 in CHARGE syndrome pathogenesis is also supported by a recent study describing 21 individuals who bear germline heterozygous mutations in *AGO2* (Lessel et al., 2020). These individuals present a complex phenotype, with many of the most frequently encountered clinical findings overlapping CHARGE syndrome (*e*.*g*., intellectual disability/developmental delay with autistic features, muscular hypotonia, visual impairment, breathing and swallowing difficulties, malformed external ear, and heart anomalies) (Hale et al., 2016).

Collectively, our data further provide evidence that CHARGE syndrome-like disease in *Ago2* mutant mice is due to the loss of a non-canonical function of AGO2, not shared with other AGO members. As demonstrated by our luciferase assays with the psi-CHECK2-let-7 × 8 reporter (Fig.5H), the observed phenotypes cannot be explained by impaired post-transcriptional gene silencing – for which AGO2 is functionally redundant with other AGO members (Su et al., 2009). Impairment of post-transcriptional gene silencing in NCCs via conditional *Dicer* deletion results in a much more severe phenotype, with an almost complete lost of craniofacial structures (Zehir, Hua et al., 2010). Our data indicate that the observed phenotypes are instead due to perturbation of cotranscriptional alternative splicing (Figs.6-7), a process that has been shown to be regulated by AGO2 in a variety of other contexts (Ameyar-Zazoua et al., 2012, Batsche & Ameyar-Zazoua, 2015, Chu et al., 2021, Kalantari et al., 2016, Taliaferro et al., 2013, Tarallo et al., 2017). Interestingly, a study in adult flies shows that the splicing regulatory function of AGO2 is dependent of its RNA-binding activity but independent of its endonuclease activity (Taliaferro et al., 2013), reminiscent of the situation described during early development of mouse embryos (Ngondo et al., 2018). Yet, the exact molecular mechanism by which AGO2 regulates alternative splicing, and the role of FAM172A in this process, are not fully understood. Both AGO2 (Ameyar-Zazoua et al., 2012) and FAM172A (Belanger et al., 2018, Belanger et al., 2022) are known to interact with many chromatin- and spliceosome-associated proteins, suggesting a role in bridging both machineries (Batsche & Ameyar-Zazoua, 2015, Berube-Simard & Pilon, 2018). Our rapamycin data are consistent with this possibility (Fig.7), further suggesting that it is possible to circumvent the loss of either AGO2 or FAM172A by increasing the pool of available spliceosome-associated proteins. Other data suggest that AGO2 may influence splicing output by reducing the elongation rate of RNAP II via small RNA-mediated interaction with antisense transcripts, thereby enhancing the inclusion of alternative exons (Ameyar-Zazoua et al., 2012). However, such a model cannot explain why some alternatively spliced exons are more included than normal when AGO2 is absent (Figs.6-7). Similarly, other RISC-associated proteins (*e*.*g*., TNRC6 and DCR) may also be involved (Ameyar-Zazoua et al., 2012, Johnson, Chu et al., 2021), but their contribution appears not to be systematic (Liu, Liu et al., 2018, Taliaferro et al., 2013). These observations thus suggest that AGO2 regulates alternative splicing through different means. Determining whether FAM172A plays a role in all or only a subset of them will require further studies.

## Materials and Methods

### Animals

All experimental procedures involving mice were approved by the institutional ethics committee of the Université du Québec à Montréal (CIPA protocols #650 and #933) in accordance with the biomedical research guidelines of the Canadian Council of Animal Care (CCAC). *Fam172a*^*Tp/+*^ (FVB/N background; also known as *Toupee*^*Tg/+*^) (Belanger et al., 2018), *Ago2*^*Flox/+*^ (C57BL/6 background; also known as *Ago2*^*tm1*.*1Tara/J*^) (O’Carroll et al., 2007), *Wnt1-Cre2*^*Tg/+*^ (C57BL/6 background; also known as *E2f1*^*Tg(Wnt1-cre)2Sor/J*^) (Lewis et al., 2013) and *Rosa26*^*FloxSTOP-YFP*^ (FVB/N background; also known as *Gt(ROSA)26Sortm1(EYFP)Cos*) (Srinivas, Watanabe et al., 2001) mouse lines were as previously described. *Ago2*^*Flox/+*^ and *Wnt1-Cre2*^*Tg/+*^ lines were purchased from The Jackson Laboratory (JAX stock #016520 and #022501, respectively), while the *Rosa26*^*FloxSTOP-YFP*^ line was directly provided by Dr. Frank Costantini (Columbia University). Embryos were generated by natural mating and staged by considering noon of the day of vaginal plug detection as e0.5. Control (WT or either *Ago2*^*Flox/+*^ or *Wnt1-Cre2*^*Tg/+*^ single transgenics) and mutant embryos were obtained from the same litter and processed in parallel. Primers used for genotyping are listed in Table S2.

For rapamycin treatments, pregnant dams of relevant genotype were injected intraperitoneally with a 165mg/ml solution (200µM; in 20% ethanol) at a final concentration of either 0.5mg or 1mg per kg of body weight. Control animals were administered 20% ethanol only. Dams were treated either once a day (around noon) or twice a day (in morning and evening, at 12h intervals) for a period of 3 days between e9.5 and e11.5, and embryos were collected at e12.5 for analysis.

### DNA constructs and recombinant proteins

MYC- and MBP-tagged versions of the 371aa-long isoform of murine FAM172A (WT and E229Q-mutated) and both psiCHECK2-let-7X8 reporter and empty psiCHECK2 vector (kindly provided by Dr. M. Simard, Université Laval) were as previously described (Belanger et al., 2018, Johnston et al., 2010). Vectors containing human *AGO1* (pcDNAM-human eIF2C1-WT-Myc; gift from Dr. K. Ui-Tei) (Doi, Zenno et al., 2003) and *AGO2* ORFs (p3XFLAG-myc-CMV-AGO2, gift from Dr. E. Chan) (Lian, Li et al., 2009) were obtained from Addgene (#50360 and #21538, respectively). For BiFC assays, full-length ORFs of *Fam172* and *AGO2* were subcloned in expression vectors bearing either the N-terminal (pCDNA3-_1-173_Venus) or the C-terminal half (pCDNA3-_174-259_Venus) of Venus, which were both generously provided by Dr. S. Merabet (Institut de génomique fonctionnelle de Lyon) (Hudry, Viala et al., 2011). NLS-mutated FAM172A (R227Q, R228Q, R241Q and R242Q), phosphodead FAM172A (D219Q, D222Q), phosphomimic FAM172A (S215D, S216D, S217D, S218D and T222D), FLAG-tagged versions of both AGO1 and AGO2 as well as NLS-containing _FLAG_AGO2 were all generated using the Gibson assembly method, as previously described (Gibson, 2011, Gibson, Young et al., 2009). Briefly, relevant PCR amplicons (see Table S2 for primers list) and digested plasmid (ratio 10:1) were added to the 1X Gibson master mix (5% PEG-8000, 100 mM Tris–HCl, pH 7.5, 10 mM MgCl2, 10 mM DTT, 200 μM each of the four dNTPs, and 1 mM NAD) and then combined to 0.01U of T5 EXO, 0.06U of Phusion polymerase and 10U/μl Taq ligase for a 1h-long incubation at 50°C. All DNA constructs were verified via Sanger sequencing.

Recombinant MBP, _MBP_FAM172A and _MBP_FAM172A[E229Q] proteins were produced in BL21 bacteria via IPTG (0.3 mM) induction of pMAL-c5X constructs, purified using amylose affinity chromatography and eluted in column buffer, as previously described (Belanger et al., 2018). Recombinant His-tagged versions of AGO1 and AGO2 proteins were purchased from Sino Biological (#11225-H07B #11079-H07B, respectively).

### Cell culture, transfection and luciferase assays

Adherent murine cells were grown in EMEM (N2a, MEFs) or DMEM (NIH3T3, R1) medium supplemented with either 10% (N2a, MEFs and NIH3T3) or 20% (R1) FBS, and 1% penicillin/streptomycin (all products from Wisent). MEFs were generated from e10.5 embryo heads dissociated in EMEM containing 1.3 mg/ml Dispase II (Life Technologies #17105–041), 0.4 mg/ml collagenase (Sigma #C2674) and 0.1 mg/ml DNase I (Sigma #DN25), for 30 min at 37°C with gentle agitation. The resulting cell suspension was filtered through a 70µm cell strainer (Fisherbrand) and then plated on gelatin-coated coverslips for immunofluorescence or gelatin-coated plates for protein extraction. Human LCLs derived from a FAM172A[E228Q]-associated CHARGE family (Belanger et al., 2018) were grown in suspension in RPMI medium supplemented with 10% FBS and 1% penicillin/streptomycin (all products from Wisent). All cell cultures were maintained at 37°C and 5% CO_2_ in a humidified atmosphere.

All transfections were performed using GeneJuice reagents (Millipore-Sigma #70967) in accordance with the manufacturer’s instructions (0.1 µg of DNA per cm^2^ of cultured cells, and 1µg of DNA for 3µl of GeneJuice reagent). For luciferase assays, heads of WT and *Ago2*-mutant e10.5 embryos also expressing the *Rosa26*^*FloxSTOP-YFP*^ reporter in NCCs were dissociated as described for MEFs and subjected to FACS for specifically recovering NCCs. After 48h transfection with appropriate plasmid (psiCHECK2-let-7X8 or empty psiCHECK2 vector) in 96-well plates, cell content was release with lysis buffer (100mM Tris pH8.0, 1% NP-40%, 1mM DTT) and luciferase activity was measured using a Lumat LB9507 luminometer (Berthold) in luciferase assay buffer (1M glycyl glycine, 1M MgSO4, 1M KH2PO4, 0.2M EGTA, luciferine).

### Immunofluorescence analysis of cultured cells

Cells plated on coverslips were transfected with the appropriate plasmid for 48h, fixed in 4% paraformaldehyde (diluted in PBS) for 15 minutes on ice and washed three times in PBS. Cells were then incubated in blocking solution (10% FBS and 0.1% TritonX-100 diluted in PBS) for 1h at room temperature, followed by overnight incubation at 4°C with primary antibodies of interest diluted in blocking solution (see Table S3 for antibodies list). After three more washes in blocking solution, cells were incubated with relevant secondary antibodies conjugated with AlexaFluor 488, 594 or 647 for 1h at room temperature (see Table S3 for antibodies list), washed again three times and finally counterstained with 5µg/ml DAPI (40,6-diamidino-2-phenylindole) for 10 min. Images were acquired using a Nikon A1 laser scanning confocal microscope with Plan Apo λ 60×1.40 objective, and analyzed using ImageJ. Mean fluorescence intensity in the nucleus and the cytoplasm was quantified using the *Measure* function and these values were used to evaluate the relative abundance of proteins of interest in each cell compartment, which was expressed in log2 scale (log2[N:C] ratio).

### Bimolecular Fluorescence Complementation (BiFC) assays in cells

Efficiency of BiFC assays in 6-well plates was assessed 24h post-transfection by calculating the percentage of Venus(YFP)-positive cells and mean fluorescence intensity using a BD Accuri C6 flow cytometer (BD Biosciences). After reaching the desired number of detected events (30,000 or 100,000 cells), remaining cells were used to verify abundance of N-Venus-and C-Venus-tagged proteins by anti-GFP western blot (see below). Protein abundance was then used for normalization of fluorescence intensity, which also took into account auto-fluorescence in non-transfected cells as background value to be subtracted for all samples. Subcellular distribution of BiFC signal was assessed 48h post-transfection in unfixed cells plated in 35mm µ-dishes (Ibidi #81156), after 5-min incubation with 5μg/ml Hoechst 33342. For live imaging, cells were plated on 8-well chamber µ-slides (Ibidi #80826), exposed to either 2ng/ml Leptomycin B (LMB) (Cell Signaling Technology #9676) or vehicle only (same volume of ethanol 100%), and monitored for an additional 5h under normal growing conditions, with image acquisition every 20 min. All images were acquired using a Nikon A1 confocal microscope and analyzed using ImageJ, as described above.

### Co-Immunoprecipitation in cells and *in vitro*

Co-IP assays in cells were performed using two confluent 100mm plates, 48h after transfection with relevant expression vectors. Whole cell extracts were prepared in previously described low-salt buffer (20 mM Tris pH 8.0, 25mM NaCl, 1 mM EGTA, 1.5 mM MgCl2, 1 mM DTT, 1% Triton TX-100, 10% Glycerol, 1X Roche Complete protease inhibitors) during 1h at 4°C, after which samples were centrifuged to remove insoluble material (Beland, Pilon et al., 2004). *In vitro* co-IP assays were performed using 300ng of purified recombinant proteins diluted in same low-salt buffer. For both cellular and *in vitro* assays, 10% of lysate was kept aside as input material and the rest was incubated with primary antibody of interest overnight at 4°C (see Table S3 for antibodies list). Antibody-bound complexes were then isolated using protein G-coupled magnetic beads (Dynabeads protein G, ThermoFisher) and analyzed either by western blot (see below) or mass spectrometry (for assessing FAM172A phosphorylation) (Belanger et al., 2018).

### Western Blot

Immunoprecipitated protein complexes or either whole cell extracts prepared in low-salt buffer (Beland et al., 2004) or cytoplasmic/nuclear fractions prepared via the REAP method (Suzuki, Bose et al., 2010) were separated on SDS-PAGE mini-gels using the Mini-Protean system (Bio-Rad Laboratories) and then transferred to PVDF membranes. Membranes were blocked at room temperature for 1h in TBST (Tris-buffered saline-Tween 20) containing 5% non-fat milk, followed by overnight incubation at 4°C with primary antibody of interest (see Table S3 for antibodies list). After three washes with TBST, membranes were subsequently incubated with relevant HRP-conjugated secondary antibodies for 1h at room temperature (see Table S3 for antibodies list). Chemiluminescent detection was finally performed using Immobilon Western HRP substrate (Millipore-Sigma) and captured using the Fusion FX imaging system (Vilber). When necessary, relative protein levels were quantified using the *Gel Analyzer* function in ImageJ.

### Embryonic and neonatal tissue analysis

Anti-βIII-Tubulin immunofluorescence, hematoxylin & eosin staining and neurofilament immunohistochemistry (see Table S3 for antibodies list) were performed on embryos fixed in 4% paraformaldehyde overnight, as previously described (Bergeron, Cardinal et al., 2015, Boulende Sab, Bouchard et al., 2011, Sanchez-Ferras, Bernas et al., 2016). Anti-βIII-Tubulin immunofluorescence and hematoxylin & eosin staining were performed on 10μm sections of OCT-embedded and paraffin-embedded tissues, respectively. Neurofilament immunohistochemistry was performed on whole embryos in PBSMT (2% milk, 0.5% triton in PBS), after bleaching with methanol/DMSO/H2O2 (4:1:1), with HRP-conjugated secondary antibody finally detected using 3,3’-diaminobenzidine tetrahydrochloride (Millipore-Sigma). For coloboma analysis, unfixed embryos were directly analyzed under brightfield light. Images were acquired using either a Leica DFC 495 camera (for colored images) or Lumenera Infinity 2.0 camera (for brightfield images) mounted on a Leica M205 FA stereomicroscope. Choroidal fissure width and organ/body cavity surface ratio were all determined using ImageJ.

### RNA extraction and RT-qPCR

RNA from e12.5 embryonic heads was extracted using the RNeasy Plus Purification Kit (Qiagen) in accordance with the manufacturer’s protocol. cDNAs were then generated using 50 ng of total RNA and Superscript II reverse transcriptase (ThermoFischer Scientific). qPCR experiments were finally carried out using the Ssofast EvaGreen Supermix and C1000 Touch thermal cycler (Bio-Rad Laboratories), with *Psmb2* gene used for normalization of absolute expression levels and constant exons of selected genes used for normalization of splicing events (see Table S2 for primers list).

### Statistical analysis

Graphical data are represented as the mean ± SD, with the number of independent experiments (N) and/or independent biological replicates (n) specified in the figure and/or legend when relevant. Significant differences between samples were determined using the GraphPad Prism software version 6.0, with 0.05 as statistical significance cut-off. Selected statistical tests are indicated in figure legends.

## Acknowledgements

The authors thank the Cellular analyses and Imaging core (CERMO-FC, UQAM) for assistance with confocal imaging and flow cytometry; the Proteomics Discovery Platform (IRCM) for the mass spectrometry analyses; Dr. S. Merabet and J. Reboulet for providing BiFC constructs and technical advice; Dr. M. Simard (Université Laval) for providing the psi-CHECK2-let-7×8 and relevant empty vectors; Dr F. Costantini (Columbia University) for providing *Rosa26*^*FloxSTOP-YFP*^ mice; and Dr. J. W. Belmont and P. Hernandez (Baylor College of Medicine) for providing the human lymphoblastoid cell lines. This work was supported by a grant from the Canadian Institutes of Health Research to NP (grant #PJT-152933). NP was also supported by the Fonds de la recherche du Québec – Santé (FRQS Senior Research Scholar) and by the UQAM Research Chair on Rare Genetic Diseases, while EL was supported by a doctoral scholarship from the Natural Sciences and Engineering Research Council (NSERC), CB was supported by a doctoral scholarship from the FRQS, and both SS and FA were supported by scholarships from the CERMO-FC.

## Author Contributions

NP conceived and supervised the study; SS, FABS and NP designed the experiments; SS, FABS, BG, EL, FA and CB performed the experiments and collected data; All authors analyzed and interpreted data; SS, FABS and NP drafted the manuscript; NP edited the manuscript; All authors revised the manuscript.

