## Supplementary material for "FAM172A controls the nuclear import and alternative splicing function of AGO2": Fig.S

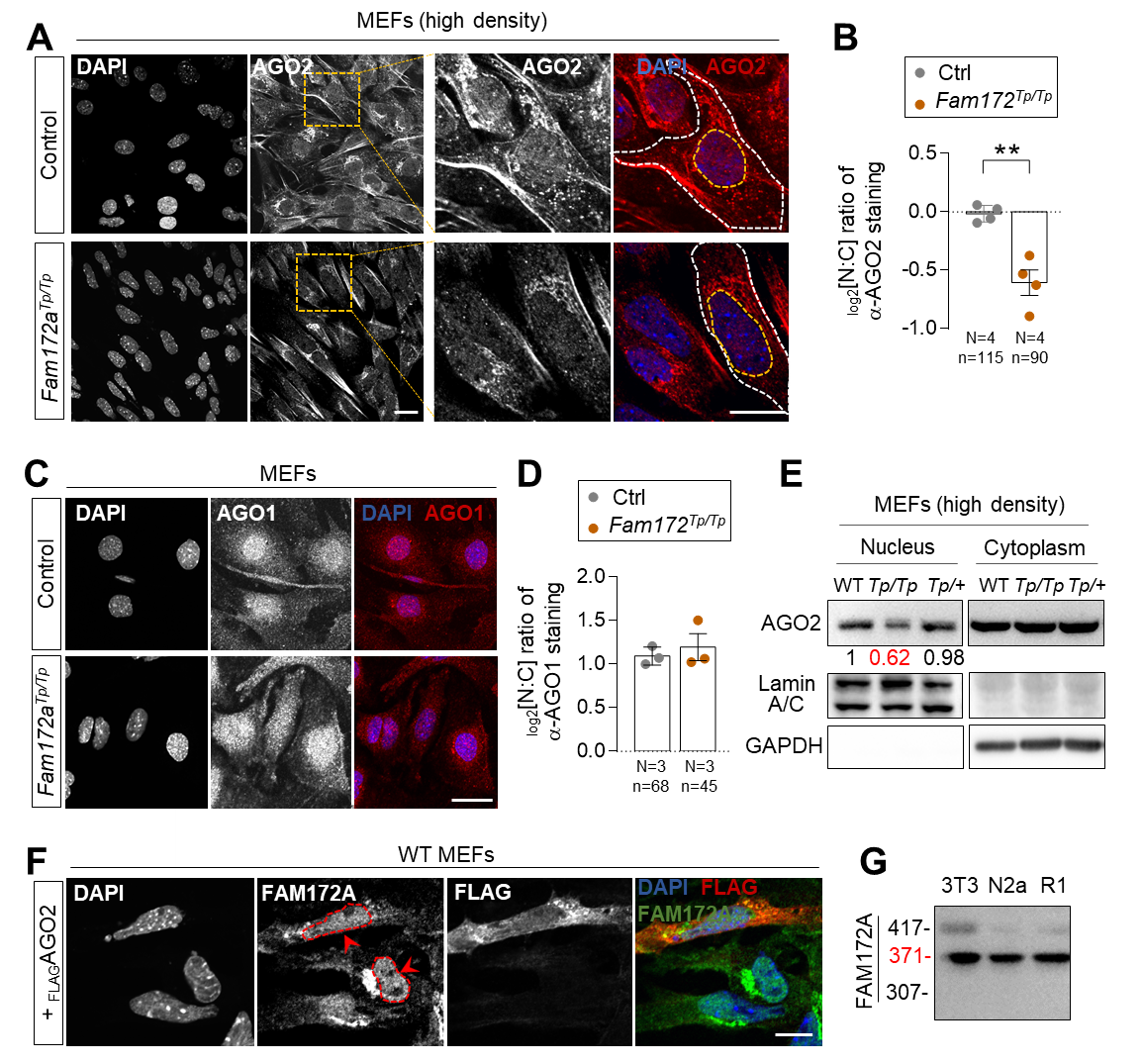
**Figure S1. FAM172A influences the nuclear localization of AGO2, but not AGO1. (A)** Immunofluorescence analysis of AGO2 distribution in WT and *Fam172^Tp/Tp^* e10.5 MEFs at high density (200 000 cells/cm^2^), with nuclei stained using DAPI. **(B)** Quantification of relative fluorescence intensity for anti-AGO2 staining in nucleus and cytoplasm (N:C ratio, expressed in log 2 scale) using images such as those displayed in A. Yellow and white dashed lines in zoomed-in views in A delineate areas of measurement for nucleus and cytoplasm, respectively. **(C)** Immunofluorescence analysis of AGO1 distribution in WT and *Fam172^Tp/Tp^* e10.5 MEFs at high density (200 000 cells/cm^2^), with nuclei stained using DAPI. **(D)** Quantification of relative fluorescence intensity for anti-AGO1 staining in nucleus and cytoplasm (N:C ratio, expressed in log 2 scale) using images such as those displayed in C. **(E)** Western blot analysis of AGO2 protein levels in nuclear and cytoplasmic fractions of WT, *Fam172^Tp/+^* and *Fam172^Tp/Tp^* e10.5 MEFs (at high density) showing a specific decrease in the nucleus of *Fam172^Tp/Tp^* cells (N=3). Lamins A/C and GAPDH are used as loading control for nuclear and cytoplasmic fractions, respectively. Indicated numbers correspond to relative amounts of nuclear AGO2 after normalization for amount of lamins A/C, as determined via densitometry. **(F)** Immunofluorescence analysis of FAM172A distribution in WT e10.5 MEFs transfected or not with **_FLAG_**AGO2 (N=3). Red arrowheads compare fluorescence intensity in nuclei of transfected and non-transfected cells. **(G)** Western blot analysis of FAM172A isoforms in NIH 3T3 fibroblasts, Neuro2a (N2a) neuroblasts, and R1 embryonic stem cells (N=3). Scale bar, 20μm. ***P* ≤ 0.01; Student’s *t*-test.

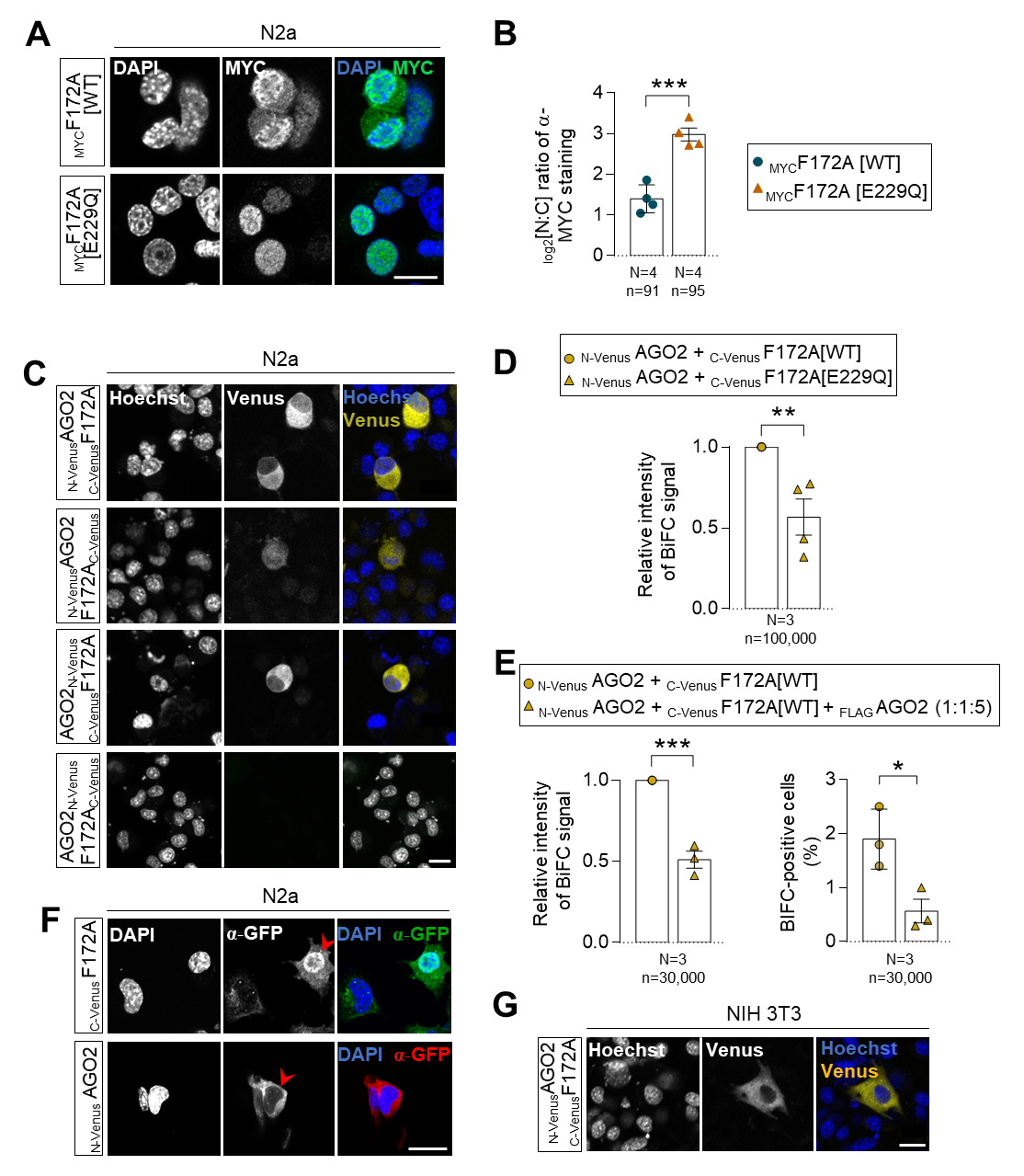
**Figure S2. FAM172A-AGO2 interaction is direct and specific. (A)** Immunofluorescence analysis of the subcellular distribution of WT and E229Q-mutated **_MYC_**FAM172A proteins in transfected N2a cells, with nuclei stained using DAPI. **(B)** Quantification of relative fluorescence intensity for anti-MYC staining in nucleus and cytoplasm (N:C ratio, expressed in log 2 scale) using images such as those displayed in A. **(C)** Distribution of AGO2-FAM172A BiFC signal in living N2a cells 48h after transfection with indicated combinations of N-Venus-tagged AGO2 and C-Venus-tagged FAM172A constructs, with nuclei stained using Hoechst (N=3). **(D-E)** Comparison of overall BiFC fluorescence intensity between _N-Venus_AGO2–_C-Venus_F172A[WT] and _N-Venus_AGO2–_C-Venus_F172A[E229Q] (D) or after competition with a 5-time excess of _FLAG_AGO2 (E; also displaying percentage of positive cells in right panel). Mean fluorescence intensity and percentage of positive cells were determined using flow cytometry 24h after transfection in N2a cells. **(F)** Immunofluorescence analysis of _C-Venus_FAM172A and _N-Venus_AGO2 distribution in transfected N2a using a GFP antibody that recognize both halves of Venus, with nuclei stained using DAPI (N=3). Red arrowheads point to transfected cells. **(G)** Distribution of _N-Venus_AGO2–_C-Venus_F172A BiFC signal in living NIH3T3 cells 48h after transfection, with nuclei stained using Hoechst (N=3). Scale bar, 20μm. **P* ≤ 0.05, ***P* ≤ 0.01 and ****P* ≤ 0.001; Student’s *t*-test.

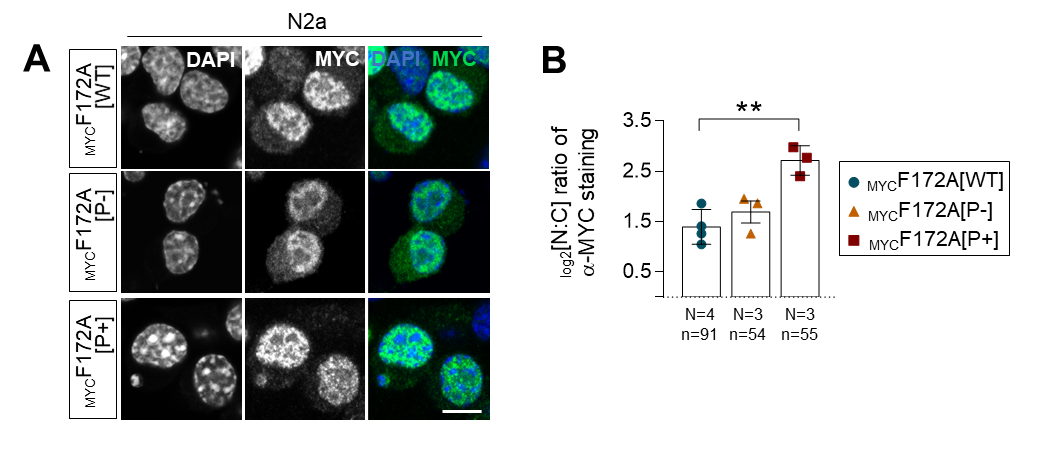

**Figure S3. Status of FAM172A phosphorylation influences its own nuclear localization. (A)** Immunofluorescence analysis of the distribution of MYC-tagged WT, phosphodead (P-) and phosphomimetic (P+) versions of murine FAM172A protein in transfected N2a cells, with nuclei stained using DAPI. **(B)** Quantification of relative fluorescence intensity of anti-MYC staining in nucleus and cytoplasm (N:C ratio, expressed in log 2 scale) using images such as those displayed in A. Data for _MYC_F172A[WT] condition are also shown in Figure S2A, being duplicated here for comparison purposes only. Scale bar, 10μm. ***P* ≤ 0.01; Student’s *t*-test.

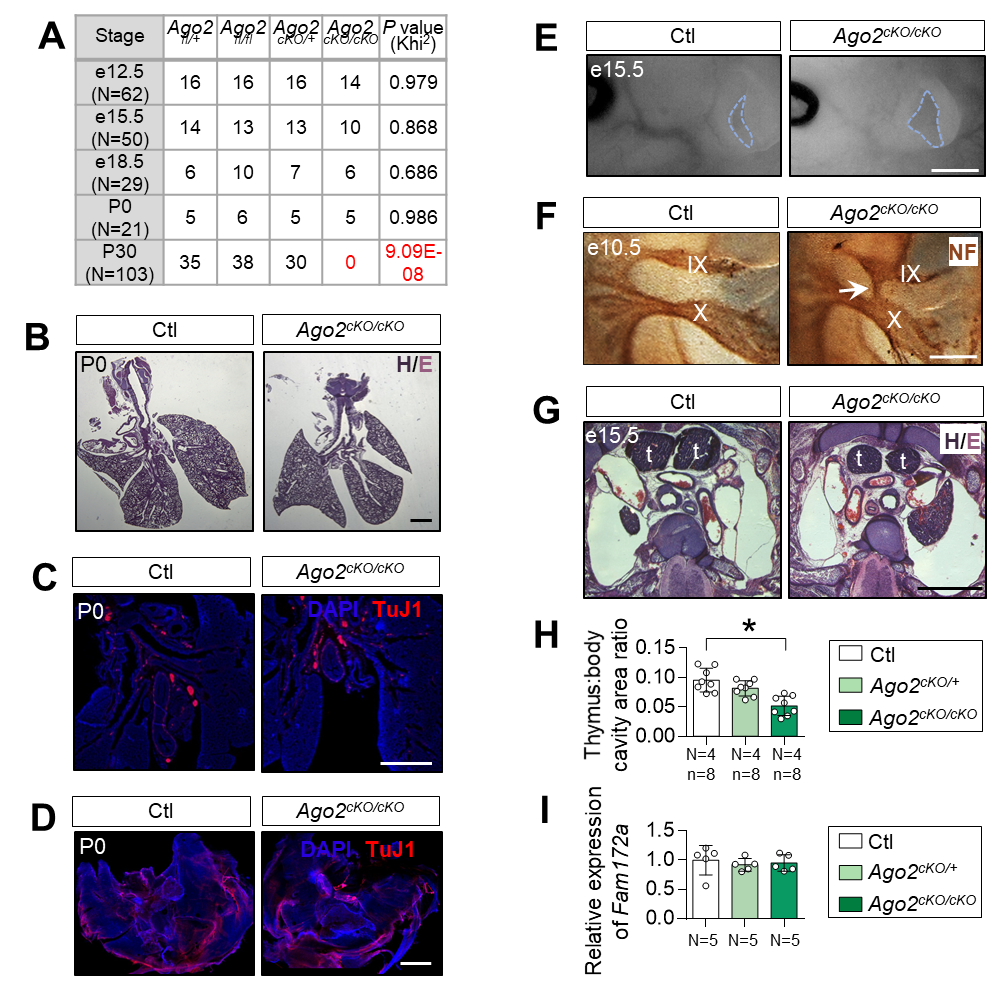

**Figure S4. NCC-specific *Ago2* knockout phenocopies CHARGE syndrome in mice. (A)** Table showing the number of animals recovered from *Wnt1-Cre2^Tg/+^*;*Ago2^Flox/+^* X *Ago2^Flox/Flox^* crosses as a function of indicated stages and *Ago2* genotypes. **(B)** H&E-stained section of P0 lungs (N=3 per genotype). **(C-D)** Immunofluorescence analysis of neuronal class III β-Tubulin (TuJ1) distribution in sections of P0 lungs (C) and whole diaphragms (D), with nuclei stained using DAPI (N=3 per genotype). **(E)** Bright-field views of e15.5 embryos, with dashed lines delineating the ear openings (N>7 per genotype). **(F)** Whole-mount staining of cranial nerves in e10.5 embryos using anti-neurofilament immunohistochemistry, with arrow pointing to an abnormal connection between glossopharyngeal (IX) and vagal (X) nerves. **(G)** H&E-stained cross-sections of e15.5 embryos at the level of thymus (t). **(H)** Quantitative analysis of thymus:body cavity area ratio, using images such as those displayed in G. **(I)** *Psmb2*-normalized RT-qPCR analysis of *Fam172a* expression in e12.5 embryo heads from *Ago2*-mutant embryos. Scale bar, 1mm. **P* ≤ 0.05; Student’s *t*-test.

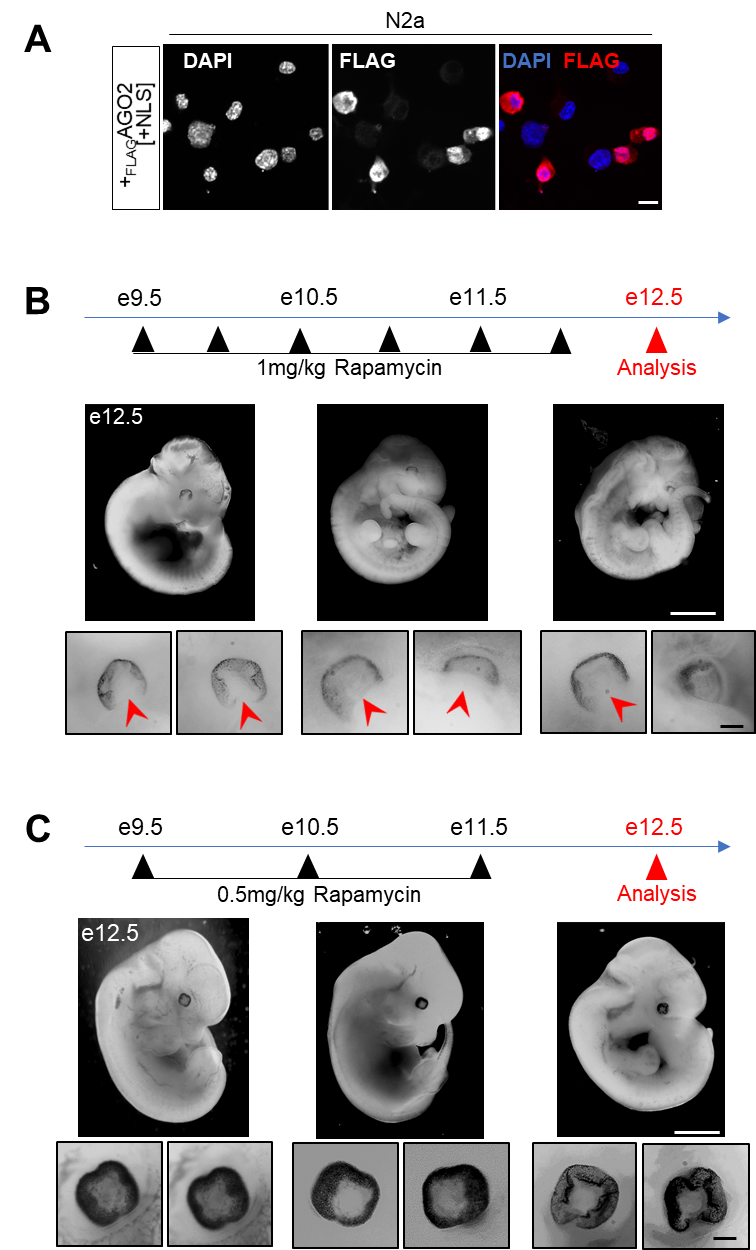

**Figure S5. Impact of acute rapamycin treatment on *Ago2*-mutant embryos. (A)** Immunofluorescence analysis of the distribution of NLS-containing _FLAG_AGO2 in transfected N2a cells, with nuclei stained using DAPI. **(B-C)** Comparison of former (B; previously used for *Fam172a^Tp/Tp^* embryos) and revised (C) treatment strategy. Rapamycin was administered to pregnant dams between e9.5 to e11.5, either at 1mg/kg body weight twice a day (B) or 0.5mg/kg body weight once a day (C). Embryos were then collected at e12.5 and brightfield images were taken to analyze gross morphology and presence/absence of coloboma (red arrowheads). Scale bar, 500μm (embryos) and 50μm (eyes).

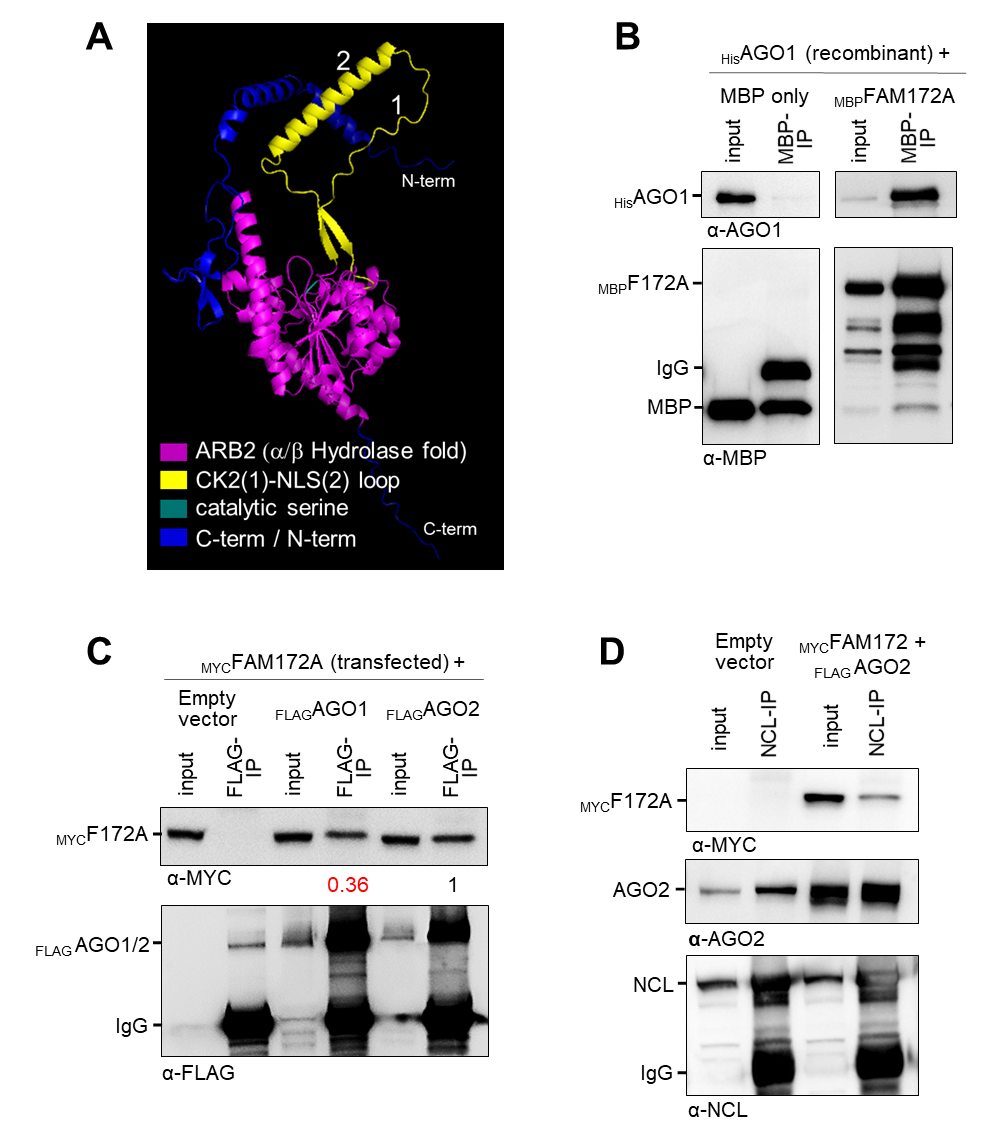

**Figure S6. FAM172A also interacts with AGO1 and NCL. (A)** PyMOL vizualisation of AlphaFold-predicted 3D structure of FAM172A. (**B**) *In vitro* co-IP of recombinant _MBP_FAM172A and _His_AGO1, using MBP as bait (N=3). MBP alone was used as negative control. **(C)** Co-IP of transfected _MYC_FAM172A and either _FLAG_AGO1 or _FLAG_AGO2 in N2a cells, using FLAG as bait (N=3). Numbers between immunoblots refer to the amount of co-immunoprecipitated _MYC_FAM172A after normalization for amount of immunoprecipitated _FLAG_AGO proteins, as determined via densitometry. **(D)** Co-IP of transfected _MYC_FAM172A and _FLAG_AGO2 with endogenous Nucleolin (NCL) in N2a cells, using NCL as bait (N=3).

**Table S1: Phenotypic overview of single Ago2^cKO/cKO^ and double Ago2^cKO/cKO^;Fam172a^Tp/+^ mutants**

| **Genotype Phenotype** | | ***Ago2^cKO/cKO^*** | ***Ago2^cKO/cKO^*;*Fam172^Tp/+^*** |
| --- | --- | --- | --- |
| **Choanal atresia** | | 100%  (n=5) | 100%  (n=5) |
| **Cleft palate** | | 0%  (n=5) | 100%  (n=5) |
|  | **Overall** | 88%  (n=8) | 69%  (n=13) |
| **Coloboma** | **Unilateral** | 86%  (n=7) | 11%  (n=9) |
|  | **Bilateral** | 14%  (n=7) | 89%  (n=9) |
|  | **+Microphtalmia** | 14%  (n=7) | 44%  (n=9) |
| **Growth retardation** | | 50%  (n=6) | 100%  (n=5) |
| **External ear malformation** | | 29%  (n=7) | N.D. |
| **Thymus hypoplasia** | | 100%  (n=4) | N.D. |
| **Cranial nerve anomalies** | | 57%  (n=7) | N.D. |

***Note****: % refers to frequency of observed phenotype; n number refers to numbers of animals; N.D., not determined.*

**Table S2: List of primers used in this study**

|  | **Oligos Name** | **Sens** | **Sequence 5’-3’** |
| --- | --- | --- | --- |
| **Genotyping** | *Fam172a^Tp^* | F  R | 5’-GGGAGTAAGTCCTACCAATGTTAAATC  5’-TGACCTTCTAAACAGTCCCATATCCCC |
|  | *Wild type* | F  R | 5’-GAAGTGGGAACAAAACACCCTTGG  5’-GTAAGGCTCGCACTGACATAGC |
|  | *Ago2^flox^* | F  R | 5’-TGATCATGGTTGAGGTCTGA  5’-GTGAGCCACTCACTGTGCAC |
|  | *Wnt1-Cre2^Tg^* | F  R | 5’-CAGCGCCGCAACTATAAGAG  5’-CATCGACCGGTAATGCAG |
|  | *Rosa26^FloxSTOP-YFP^* | F  R | 5’-CGACCTGCAGGTCCTCG  5’-CTCGAGTTTGTCCAATTATGTCAC |
| **Mutagenesis** | Gibson for FAM172A in backbone pCDNA3.1 | N-term-F  C-term-R | 5’-CTCGGCATGGACGAGCTGTACAAGGAATTCATGAAAAAGGACGAACCACCTT  TT  5’-CACTATAGAATAGGGCCCTCTAGACTCACAGCTCCTCGTGC |
|  | Gibson for FAM172A in backbone pIRES2-GFP | N-term-F  C-term-R | 5’-TTTCTGAAGAAGATCTGAGATCTCTCGAGATGAAAAAGGACGAACCACCTTTT  5’-GGATCCCGCGGGTCGACCTGGAGGTCACAGCTCCTCGTGC |
|  | FAM172A [mNLS] | N-term-R  mNLS-F  mNLS-R  C-term-F | 5’-CCTTTCCTGTTGCCCTGCTGGCTCATCTG  5’-CAACAGGAAAGGAGAGATAAGGTCTCCAAGGAAACAAAGAAGCAACAGGAT  TTCATGAGAAG  5’-ATCCTGTTGCTTCTTTGTTTCCTTGGAGACCTTATCTCTCCTTTCCTGTTGC  CCTGTGGCTC  5’-AAGCAACAGGATTTCTATGAGAAGTACCGCAACCC |
|  | FAM172A[P-] | N-term-R  P [-] -F  P [-] -R  C-term-F | 5’-ATGCATTTTCTGCTTTTCCACTTCTATATAGTTTTCATTTGGGTT  5’-GAAGTGGAAAAGCAGAAAATGCATAAACAGTCATCATCTTCTCAAGGTACACA  GGAGCCAGCAGGGAAG  5’-TTCCCGCTTCCCTGCTGGCTCCTGTGTACCTTGAGAAGATGATGACTGTTTAT  GCATTTTCTGCTTTTCCACTTC  5’-GATGAGCCAGCAGGGAAGCGGGAAAGGAGAGATAAG |
|  | FAM172A[P+] | N-term-R  P [+] -F  P [+] -R  C-term-F | 5’-ATGCATTTTCTGCTTTTCCACTTCTATATAGTTTTCATTTGGGTT  5’-GAAGTGGAAAAGCAGAAAATGCATAAACAGGACGATGATGACGATGGTGACG  ATGAGCCAGCAGGGAAG  5’-TTCCCGCTTCCCTGCTGGCTCATCGTCACCATCGTCATCATCGTCCTGTTTAT  GCATTTTCTGCTTTTCCACTTC  5’-GATGAGCCAGCAGGGAAGCGGGAAAGGAGAGATAAG |
|  | _FLAG_AGO1[WT] | 3xFLAG-F  3xFLAG-R  Ago1-F  Ago1-R | 5’-TACGACTCACTATAGGGAGACCCAATGGACTACAAAGACCATGACG  5’-GGGTCCCGCTTCCTTGTCATCGTCATCCTTGTAATC  5’-GACGATGACAAGGAAGCGGGACCCTCG  5’-GGTGACACTATAGAATAGGGCCCTTCAAGCGAAGTACATGGTGC |
|  | _FLAG_AGO2[WT] | 3xFLAG-F  3xFLAG-R  Ago2-F  Ago2-R | 5’-TACGACTCACTATAGGGAGACCCAATGGACTACAAAGACCATGACG  5’-CTGGAATGGGTGCTTGTCATCGTCATCCTTGTAATC  5’-GACGATGACAAGCACCCATTCCAGTGGTGTAAC  5’-CTTTTTTGGATCAGCAAAGTACATGGTGCGC |
|  | _FLAG_AGO2[+NLS] | +NLS-F  +NLS-R | 5’-ATGTACTTTGCTGGTGGTGATCCAAAAAAGAAGAGAAAGGTAGATCCAAAAA  AGAAGAGAAAGGTAGATCCAAAAAAGAAGAGAAAGGTATGAAGGGCCCTATTCTATAGTGTCACC  5’-GGTGACACTATAGAATAGGGCCCTTCATACCTTTCTCTTCTTTTTTGGATCTAC  CTTTCTCTTCTTTTTTGGATCTACCTTTCTCTTCTTTTTTGGATCACCACCAGCAAAGTACAT |
| **qPCR** | *Cd44*_vE8-9 | F  R | 5’-TACCCCAGTTTTTCTGGATCAGG  5’-GCCATCCTGGTGGTTGTCTG |
|  | *Cd44*_vE13-14 | F  R | 5’-TGGAAGACTTGAACAGGACAGG  5’-GTTTTCGTCTTCTTCCGGCTC |
|  | *Cd44*_cE4-5 | F  R | 5’-ACAGACCTACCCAATTCCTTCG  5’-GGGTGCTCTTCTCGATGGTG |
|  | *Mical2*_vE8-9 | F  R | 5’-GGACAGTACCCACTACTTTGTC  5’-CCGAACACAGCAGCATCTCT |
|  | *Mical2*_vE18-19 | F  R | 5’-CGGGTCTCAGGCATAGGTAAG  5’-AGCCGGTACTTGGTTGAGTTC |
|  | *Mical2*_cE3-4 | F  R | 5’-GTGGCACAAACTGGATAAGCG  5’-GCAGGACATTGTTCCGGGAG |
|  | *Ift74*_vE10-11 | F  R | 5’-ACACTTCAGCAACAGCTAGATTC  5’-GGCGATCCCATGCTTTTGTC |
|  | *Ift74*_cE5-6 | F  R | 5’-TGTACAACCAAGAAAATTCAGTGT  5’-TGTTGTAGTCTGCTAGTTGTCCT |
|  | *Col5a3*_vE6-7 | F  R | 5’-CCCCAAAGACGATGAACCAG  5’-TCTCTGTCTCAGGGATGTGGA |
|  | *Col5a3*_vE28-29 | F  R | 5’-TACCTCTGGTAACCGGGGTCTC  5’-CCTTTTGGTCCCTCATCACCC |
|  | *Col5a3*_vE66-67 | F  R | 5’-GCAGCCCATCAGAGGTTCAC  5’-AAGAGGGTCTTCGCCTGTCC |
|  | *Col5a3*_cE10-11 | F  R | 5’-GGACAGCAGTTTGAGGGG  5’-CACGGTCCCCAGGGAAG |
|  | *Fam172a* | F  R | 5’-AGGTGACTGCTGTGGCATTGAC  5’-GGCTTCTGCGAGCTGCTCTT |

**Table S3: List of antibodies used in this study**

|  | **Antibody Name** | **Dilution** | **Source** | **RRID** |
| --- | --- | --- | --- | --- |
| **Primary antibodies** | Rabbit polyclonal Anti-FAM172A | WB (1:1000)  IF (1:500)  IP (2 μg/ml) | Abcam ab1213 | 64 AB_11127114 |
|  | Rabbit polyclonal Anti-AGO2 | WB (1:1000)  IF (1:200)  IP (3 μg/ml) | Abcam ab32381 | AB_867543 |
|  | Mouse monoclonal Anti-AGO2 | WB (1:1000)  IF (1:200)  IP (2μg/ml) | Abcam ab57113 | AB_2230916 |
|  | Rat monoclonal Anti-AGO1 | WB (1:1000)  IF (1:500)  IP (3 μg/ml) | Sigma SAB4200084 | AB_10602786 |
|  | Mouse monoclonal Anti-Neurofilament | IHC (1:500) | DSHB 2H3 | AB_2618380 |
|  | Mouse monoclonal Anti-βIII Tubulin | IF (1:500) | Abcam ab78078 | AB_2256751 |
|  | Rabbit polyclonal Anti-Ki67 | IF (1:1000) | Abcam ab15580 | AB_443209 |
|  | Mouse monoclonal Anti-MYC tag | WB (1:500)  IF (1:100) | In house hybridoma  (9E10) | - |
|  | Goat polyclonal Anti-MYC tag | WB (1:1000) | Abcam ab9132 | AB_307033 |
|  | Mouse monoclonal Anti-FLAG tag | WB (1:1000)  IP (2μg/ml) | Sigma F1804 | AB_262044 |
|  | Rat monoclonal Anti-MBP tag | WB (1:1000)  IP (2μg/ml) | Covance Research Products MRT-131P | AB_10720557 |
|  | Mouse monoclonal Anti-GAPDH | WB (1:3000) | Santa Cruz Biotech sc-32233 | AB_627679 |
|  | Rabbit polyclonal Anti-GFP | WB (1:3000)  IF (1:500) | Abcam ab290 | AB_303395 |
|  | Rabbit polyclonal Anti-lamin A/C | WB (1:1000) | Abcam ab227176 | - |
|  | Rabbit Polyclonal Anti-Nucleolin | WB (1:1000)  IP (2μg/ml) | Abcam ab50279 | AB_881762 |
| **Secondary antibodies** | Donkey Alexa Fluor 488 Anti-Rat IgG | IF (1:500) | Jackson ImmunoResearch  712-545-150 | AB_2340683 |
|  | Donkey Alexa Fluor 594 Anti-Rabbit IgG | IF (1:500) | Jackson ImmunoResearch  711-585-152 | AB_2340621 |
|  | Donkey Alexa Fluor 647 Anti-Mouse IgG | IF (1:500) | Jackson ImmunoResearch  715-605-150 | AB_2340862 |
|  | Goat Anti-Rat IgG HRP | WB (1:10000) | Santa Cruz Sc-2032 | AB_631755 |
|  | Horse Anti-Mouse IgG HRP | WB (1:2000) | Cell signaling 7076 | AB_330924 |
|  | Goat Anti-Rabbit IgG HRP | WB (1:500) | Abcam 6721 | AB_955447 |
|  | Rabbit Anti-Goat IgG HRP | WB (1:25000) | Bio-Rad 1721034 | AB_11125144 |

**Movie S1. Subcellular distribution of AGO2-FAM172A BiFC signal in LMB-treated N2a cells, using WT _C-Venus_FAM172A.**

**Movie S2. Subcellular distribution of AGO2-FAM172A BiFC signal in LMB-treated N2a cells, using NLS-mutated _C-Venus_FAM172A.**

**Movie S3. Subcellular distribution of AGO2-FAM172A BiFC signal in LMB-treated N2a cells, using phosphodead _C-Venus_FAM172A.**

**Movie S4. Subcellular distribution of AGO2-FAM172A BiFC signal in LMB-treated N2a cells, using phosphomimetic _C-Venus_FAM172A.**

**Movie S5. *Ago2^cKO/cKO^* newborns die from respiratory failure.**
